# ‘*Candidatus* Phytoplasma zeae’: community-driven species delineation of the maize bushy stunt phytoplasma, a *Dalbulus*-transmitted corn pathogen confined to the Americas

**DOI:** 10.1101/2025.08.28.671991

**Authors:** Thierry Alexandre Pellegrinetti, Joshua Molligan, Abraão Almeida Santos, Maria Cristina Canale, Maira Rodrigues Duffeck, Alejandro Olmedo-Velarde, Jordanne Jacques, Ivair Valmorbida, Ashleigh Faris, Marcos Vinicius Silva de Andrade, Magda Alana Pompeli Manica, Mauricio Luna, Paola Reyes-Caldas, Jairo Rodriguez-Chalarca, Tim Dumonceaux, Edel Pérez-Lopez

**Affiliations:** Département de phytologie, Faculté des sciences de l’agriculture et de l’alimentation, Université Laval, Québec City Québec, Canada; Centre de recherche et d’innovation sur les végétaux, Université Laval, Québec City Québec, Canada; Institute de Biologie Intégrative et des Systèmes, Université Laval, Québec City Québec, Canada; L’Institute EDS, Université Laval, Quebec City, Québec City, Québec, Canada; Empresa de Pesquisa Agropecuária e Extensão Rural de Santa Catarina (Epagri), Servidão Ferdinando Ricieri Tusseti, 89.804-904, São Cristóvão Chapecó, SC, Brazil; Department of Entomology and Plant Pathology, Oklahoma State University, Stillwater, OK 74078; Department of Plant Pathology, Entomology and Microbiology, Iowa State University, Ames, IA 50011 USA; Division of Plant Science and Technology, University of Missouri, Columbia, MO, USA, 65201; Facultad de Ciencias Agrícolas, Universidad Veracruzana, Lomas del Estadio s/n, CP 91000, Xalapa, Veracruz, México; Universidad de Los Andes, La Candelaria, Bogotá, Cundinamarca, Colombia; Crops for Nutrition and Health, The Alliance of Bioversity International and Centro Internacional de Agricultura Tropical (International Center for Tropical Agriculture), Palmira, Colombia; Saskatoon Research and Development Centre, Science and Technology Branch, Agriculture and Agri-Food Canada, 107 Science Place, Saskatoon, SK, S7N 0X2, Canada

## Abstract

A novel phytoplasma species, ‘*Candidatus* Phytoplasma zeae’, is proposed based on ecological distinctiveness, vector specificity, whole-genome comparisons, and community consensus. This phytoplasma is associated with maize bushy stunt (MBS) disease in corn (*Zea mays*) and is transmitted exclusively by D*albulus maidis* and *D. elimatus*, two leafhopper species endemic to the Americas, and has been reported in Brazil, Colombia, Mexico, Peru, and several U.S. states. Here we sequenced and assembled the genome of MBS phytoplasma strains from Brazil, and U.S. to describe and propose this new species. Although the 16S rRNA gene sequence of the proposed reference strain, MBSP-BR^RS^, shares >99% identity with that of ‘*Ca*. Phytoplasma asteris’, key nucleotide polymorphisms distinguish ‘*Ca*. P. zeae’ from other 16SrI-related phytoplasma species. Average nucleotide identity (ANI) and average amino acid identity (AAI) values between ‘*Ca*. P. zeae’ and ‘*Ca*. P. asteris’ are 97.70–98.00% and 96.65–96.88%, respectively, both near the established species delineation thresholds. Comparative genomic analyses revealed unique gene clusters in ‘*Ca*. P. zeae’ associated with amino acid transport, defense mechanisms, and protein turnover, which may contribute to its specialization in corn. The ecological profile of ‘*Ca*. P. zeae’, including its narrow host range and restricted geographic distribution, supports its recognition as a novel species under Rule c of the IRPCM guidelines. The designation ‘*Candidatus* Phytoplasma zeae’ is therefore proposed by members of the research community who have studied this pathogen for over a decade, with the MBSP-Brazil^RS^ strain serving as the reference.

## INTRODUCTION

Maize (*Zea mays* L.) is one of the world’s most important cereal crops, ranking third in production volume after wheat and rice. Each year, approximately 380 million tons are produced across 53 countries (http://www.fao.org/faostat), with cultivation widespread in nearly all tropical and temperate regions. In tropical and subtropical areas of the Americas, maize production is threatened by a serious disease caused by insect-transmitted bacteria of the genus ‘*Candidatus* Phytoplasma’ [1].

Maize bushy stunt (MBS) was first detected in Mexico in 1955 [2] and linked to a phytoplasma pathogen two decades later [3]. Since then, MBS phytoplasma (MBSP) has been reported throughout Central and South America [4-6], and more recently in Mexico in association with native maize varieties [8]. The phytoplasma associated with MBS disease has been classified as a ‘*Ca*. P. asteris’-related strain within the 16SrI-B subgroup [8,9]. MBSP exclusively infects corn and teosinte relatives and is transmitted solely by leafhoppers of the genus *Dalbulus* (Hemiptera: Cicadellidae), highlighting host specialization and co-evolutionary relationship among the plant, insect vector, and pathogen [7]. Typical MBS symptoms include leaf reddening and yellowing, stunted growth, and a characteristic bushy appearance (shortened internodes), impacting both commercial hybrids and native maize landraces [7, 10].

Due to their extreme difficulty in being cultured in axenic media, phytoplasmas lack measurable phenotypic characteristics for classical taxonomic classification. The International Committee of Systematic Bacteriology Subcommittee for the Taxonomy of Mollicutes, together with the International Research Program for Comparative Mycoplasmology (IRPCM), addressed this by establishing taxonomic conventions for uncultured bacteria [11] and creating the provisional genus ‘*Candidatus* Phytoplasma’ [12]. Since the first ‘*Ca*. Phytoplasma’ species was described 30 years ago [14], more than 50 species have been formally recognized [14-18]. According to IRPCM guidelines [13], three principal criteria are used to define new ‘*Ca*. Phytoplasma’ taxa: (1) a unique 16S rRNA gene sequence (≥1200 bp) defining the reference strain, with closely related strains distinguished by minor sequence differences; (2) less than 97.5% sequence identity with the 16S rRNA gene of any described species, or (3) if above this threshold, evidence that the organism represents an ecologically distinct population (“Rule c”), demonstrated by differences in insect vectors, plant hosts, and supporting molecular divergence. Recent revisions to these delineation criteria also incorporate analyses of additional housekeeping genes and whole-genome average nucleotide identity (ANI), adopting the general bacterial species boundary of >95–96% ANI [19,20].

When describing ‘*Ca*. P. asteris’ (16Sr I), Lee et al. [9] suggested that members of the 16SrI group might encompass multiple distinct species. To date, ‘*Ca*. P. asteris’, ‘*Ca*. P. lycopersici’, and more recently ‘*Ca*. P. triticii’, the latter defined under Rule c, have been formally delineated within this group [18,21]. In collaboration with colleagues from Brazil and the United States, we have now confirmed the presence of MBS phytoplasma in *D. maidis* (DeLong & Wolcott) (Hemiptera: Cicadellidae) from laboratory colonies originally established with insects collected in maize fields showing MBS symptoms in Brazil, as well as in field-collected specimens from the United States (Oklahoma and Iowa), where plants also showed MBS symptoms. These detections, represent the first confirmed reports of MBSP likely causing MBS in these U.S. states. Moreover, we successfully retrieved the near-complete genome sequence of the pathogen from each of these locations. Phylogenetic analyses based on housekeeping genes, comprehensive whole-genome ANI comparisons, together with the distinct ecology of this phytoplasma, characterized by its specific insect vector, host specialization, and geographic restriction, provide robust evidence supporting its establishment as a novel taxon. Taken together, these findings support the designation of ‘*Candidatus* Phytoplasma zeae’, with the MBSP-BR^RS^ (MBSP-BR) strain proposed as the representative of this newly described species.

### MAIZE BUSHY STUNT DISEASE: PLANT HOST AND SYMPTOMS

Maize bushy stunt (MBS) was first detected on corn plants in Mexico over six decades ago [2]. The main symptoms of MBS include chlorotic striping on the leaves, foliar reddening and yellowing, stunted growth, lateral branching along the stalk, deformed or poorly filled ears, and a characteristic bushy appearance [22,23]. These symptoms are consistently observed in both commercial hybrids and native landraces of maize [7,10] (**Figure 1A**). Crop losses attributed to MBS phytoplasma infection can range from 35% to as high as 93%, depending largely on the prevalence of the insect vectors and timing of the infection [24,25]. Although other members of the genus *Zea* have been shown experimentally to be susceptible to MBS phytoplasma, *Z. mays* remains the only confirmed natural host [10].

**Figure 1.**
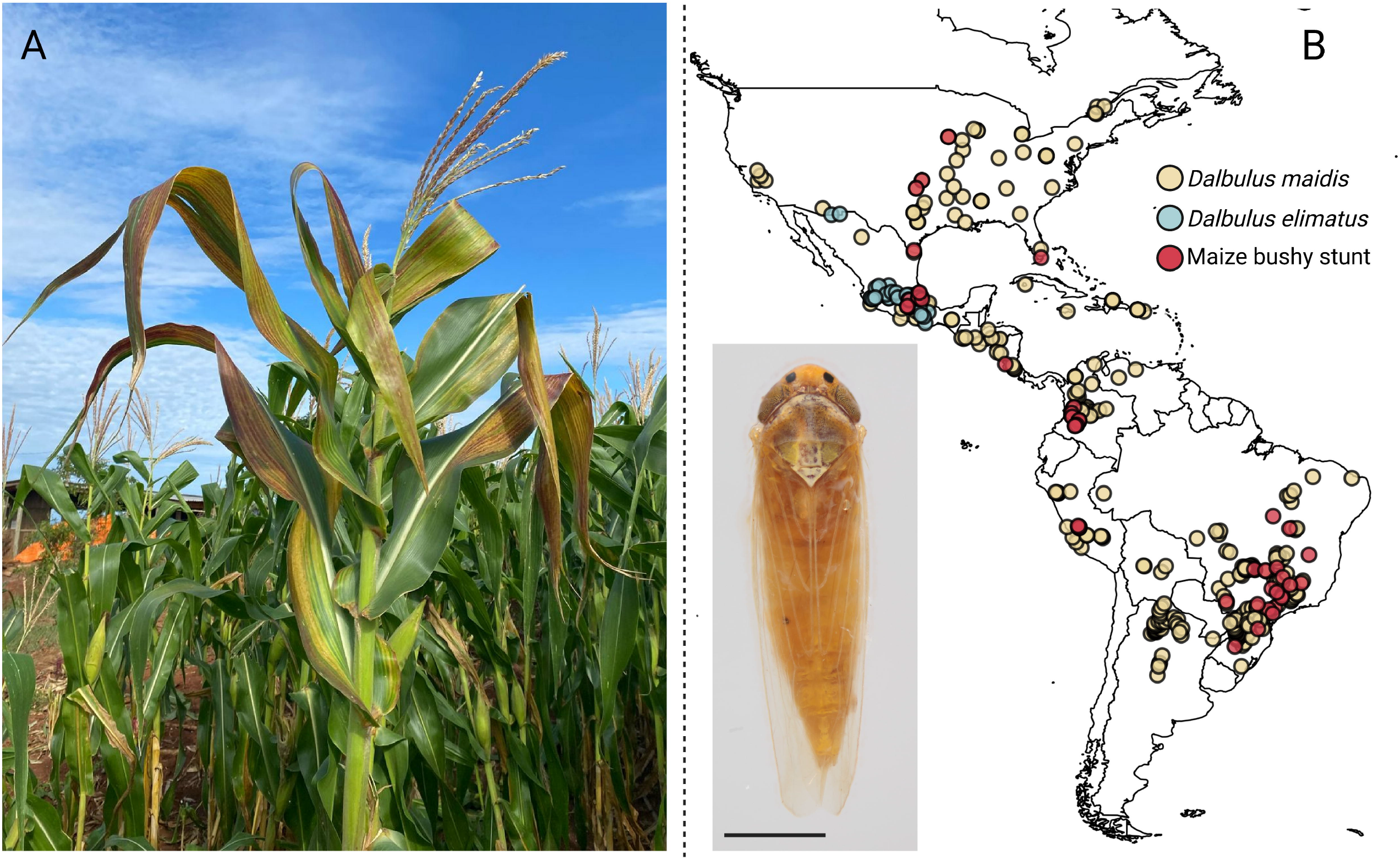
Unique host, vector and geographic distribution of Maize Bushy Stunt phytoplasma **A**) Maize plant showing typical symptoms of Maize Bushy Stunt disease, including leaf reddening and yellowing, and stunting, observed in a field in central and south Brazil. **B**) Geographic distribution of *Dalbulus maidis* (yellow), *D. elimatus* (green), and confirmed reports of Maize Bushy Stunt Phytoplasma (red) across the Americas. The inset image shows an adult *D. maidis* specimen (scale bar = 1 mm), the primary insect vector associated with MBSP transmission.

### THE GENUS DALBULUS: KEY VECTORS AND THEIR ASSOCIATION WITH ZEA SPP

*Dalbulus* (Hemiptera: Cicadellidae) is a Neotropical genus comprising eleven recognized species, all specialized in corn (*Z. mays*), related teosintes (*Zea* spp.), or *Tripsacum* spp. as host plants [26,27]. Among these, *D. maidis* and *D. elimatus* (Ball) are the most prominent, feeding on both maize and teosintes [26,27]. Other species such as *D. gelbus, D. longulus*, and *D. guevarai* are associated with maize and/or *Tripsacum* spp., whereas *D. quinquenotatus, D. guvnani, D. chiapensis, D. gramalotes, D. charlesi*, and *D. tripsacoides* are associated exclusively with *Tripsacum* spp. [27].

Although experimental studies have shown that other *Dalbulus* species and leafhoppers from different genera (e.g., *Graminella nigrifrons* Forbes) can acquire and transmit MBSP under controlled conditions, only *D. maidis*, commonly known as the corn leafhopper, and *D. elimatus*, known as the Mexican corn leafhopper, are confirmed natural vectors [10,28-30]. Recently in Brazil, the invasive planthopper *Leptodelphax maculigera* (Stål) (Hemiptera: Delphacidae) was found to be simultaneously infected with MBSP, maize rayado fino virus and maize striate mosaic virus [31, 32]. However, further laboratory experiments indicated that this species is not a vector of the MBSP, and pearl millet (*Pennisetum glaucum*) is its primary host, rather than maize [33]. These aspects highlight a high degree of vector specificity in the transmission ecology of MBSP.

### GEOGRAPHIC DISTRIBUTION OF MBSP AND ITS VECTORS

Extensive surveys and historical records indicate that MBSP and its two natural vectors, *D. maidis* and *D. elimatus*, are restricted to the Americas [6,7,34]. *D*. maidis is widely distributed from the southern United States through Central America and into South America, reaching as far south as Argentina, although the disease has not yet been reported there [26]. Of growing concern is the continued northward expansion of *D. maidis* habitats. Recent records now document its presence in southern U.S. states, including Texas and Oklahoma, as well as multiple key Midwest agricultural regions, with individuals captured in Illinois, Indiana, Iowa, Kansas, Kentucky, Michigan, Minnesota, Missouri, Nebraska, and Wisconsin in 2024 [35,36]. Remarkably, our group recently reported the first detection of *D. maidis* in Québec, Canada, representing the northernmost record to date (**Table S1**) [37]. This new occurrence could be linked to extreme weather events or shifts associated with increased maize cultivation as well as broader climate change impacts (**Figure 1B**) [38].

Today, *D. maidis* is recognized as the principal vector of ‘*Ca*. P. zeae’ and is the most prevalent leafhopper species found in maize agroecosystems across its range [27,31]. Its evolutionary expansion is thought to have occurred alongside the domestication and spread of *Z. mays* from its wild teosinte ancestors [39]. In contrast, reports of *D. elimatus* remain limited and are mainly distributed within the Nearctic region, from southern parts of the United States (Arizona and New Mexico) into Mexico. This species is believed to have originated in central Mexico and typically prefers elevations above 1,000 m (**Figure 1B**) [26,28,40,41].

### PHYLOGENETIC EVIDENCE SUPPORTING MBSP AS A NOVEL ‘CANDIDATUS PHYTOPLASMA’ TAXON

Following the detection of MBSP infecting native maize in highland communities of Veracruz, Mexico, in 2014 [7], our group continued monitoring the area, collecting symptomatic corn plants in 2015 and 2016. To characterize the MBSP strains present over these three consecutive years, we amplified the 16S rRNA operon, the *rpoB* gene, and the *cpn60* UT region from one representative sample collected each year, using previously described protocols [42-44]. The samples analyzed were MBS-Ver14 (previously identified as NMBSP-V41 [7], with existing 16S rRNA [KT444671] and *cpn60* [KT444673] sequences), MBS-Ver15, and MBS-Ver16, corresponding to collections from 2014, 2015, and 2016, respectively.

Amplified sequences generated in this study were assembled using the Staden package [45] and compared to reference sequences in GenBank via BLAST (http://www.ncbi.nlm.nih.gov). All three loci from these samples shared 99–100% nucleotide identity, including as with previously characterized MBSP strains such as MBS and MBS-Col. These results confirmed not only the continued presence of MBS phytoplasma in Veracruz across multiple years but also suggested ongoing reproduction and persistence of local vector populations. The sequences were subsequently incorporated into phylogenetic analyses and used to identify unique genetic regions within MBSP.

At that time, however, we were unable to amplify the full 16S rRNA operon or other key taxonomic markers, such as *tufB, secY, secA*, or *rplV*–*rpsC* [17], which limited resolution in strain-level comparisons. We did, however, successfully amplify the commonly used 16S F2nR2 fragment, and these partial sequences were deposited in GenBank (MBS-Ver015 – KY595947; MBS-Ver016 – KY595948). Although these sequences confirmed the presence of MBSP, they lacked sufficient phylogenetic resolution to determine the origin of the strains. Motivated by our recent detection of *D. maidis* in Québec, Canada, the northernmost record to date [37], we sought to investigate: (*i*) whether MBSP had been introduced to this new geographic area by its vector, and (*ii*) if so, whether the local strain could be traced to known populations, particularly from the U.S. and Brazil.

Genomic DNA was extracted as previously described [46], from insect and plant tissue (**Table S2**) and preparation of Illumina short-read libraries and subsequent sequencing were performed by Genome Québec (Montréal, Canada), resulting in five paired-end (PE) files of 215⍰M (BR), 44 ⍰M (QC), 113 M (OK), 282 (IA) and 171 ⍰M (MO) reads, respectively (**Table S2**). For genome reconstruction, raw reads were quality-trimmed using fastp v.0.23.4 [47]. Taxonomic classification was performed with Kraken2 v.2.1.3 [48] against the core-nt database. To eliminate host-derived reads (e.g., from leafhoppers, humans, and plants), we used KrakenTools v.1.2 (https://github.com/jenniferlu717/KrakenTools). Filtered reads were assembled using two metagenomic assemblers, MEGAHIT v.1.1.4 [49] and MetaSPAdes v.4.1.0 [50], to evaluate assembler performance and enhance contig recovery. Phytoplasma contigs were then identified by nucleotide BLAST using the MBSP-M3 genome (GCA_001712875.1) as alignment target [22]. This step enabled the recovery of near-complete genomes. Assemblies were subsequently scaffolded using RagTag v2.1.0 [51] and polished with Pilon v1.23.0 [52]. Genome integrity was measured by using completeness and contamination scores calculated with CheckM and CheckMm software [53,54]. Circular genome visualization and COG annotation were performed using GenoVi v0.2.16 [55]. Genome annotation was carried out with Bakta v1.11.3 [56], and comparative genomic analyses were conducted using the pangenome workflow implemented in Anvi’o v8 [57].

Comparative analyses revealed variation among the three reconstructed genomes in terms of sequence length, GC content, and the number of predicted coding sequences (**Figure 2A, Table S3**). All genomes were near-complete, with CheckM estimates of completeness ranging from 97.14% to 98.57% and 0% contamination. CheckM2 results showed similar high quality, with completeness values between 95.27% and 97.69%, and contamination ranging from 0.05% to 0.14%. For instance, the MBSP M3 genome available in NCBI exhibited 98.57% completeness and 0% contamination with CheckM, and 97.17% completeness and 0.06% contamination with CheckM2. Each genome consisted of two contigs, with total lengths ranging from 567,501 to 569,327 bp and containing 522 to 530 predicted coding sequences (**Figure 2A**). The genomes exhibited high integrity, with 31 to 32 tRNAs, five rRNAs, and elevated N50 values ranging from 561,808 to 564,331 bp. Functional annotation based on COG categories showed a predominance of genes involved in translation, ribosomal structure, and biogenesis, followed by those related to transcription and DNA replication, consistent with the streamlined genomes typical of phytoplasmas. Minor differences were observed in metabolic and cellular process categories (e.g., M, N, O), potentially reflecting adaptations to different hosts or environments (**Figure 2B, Table S4**).

**Figure 2.**
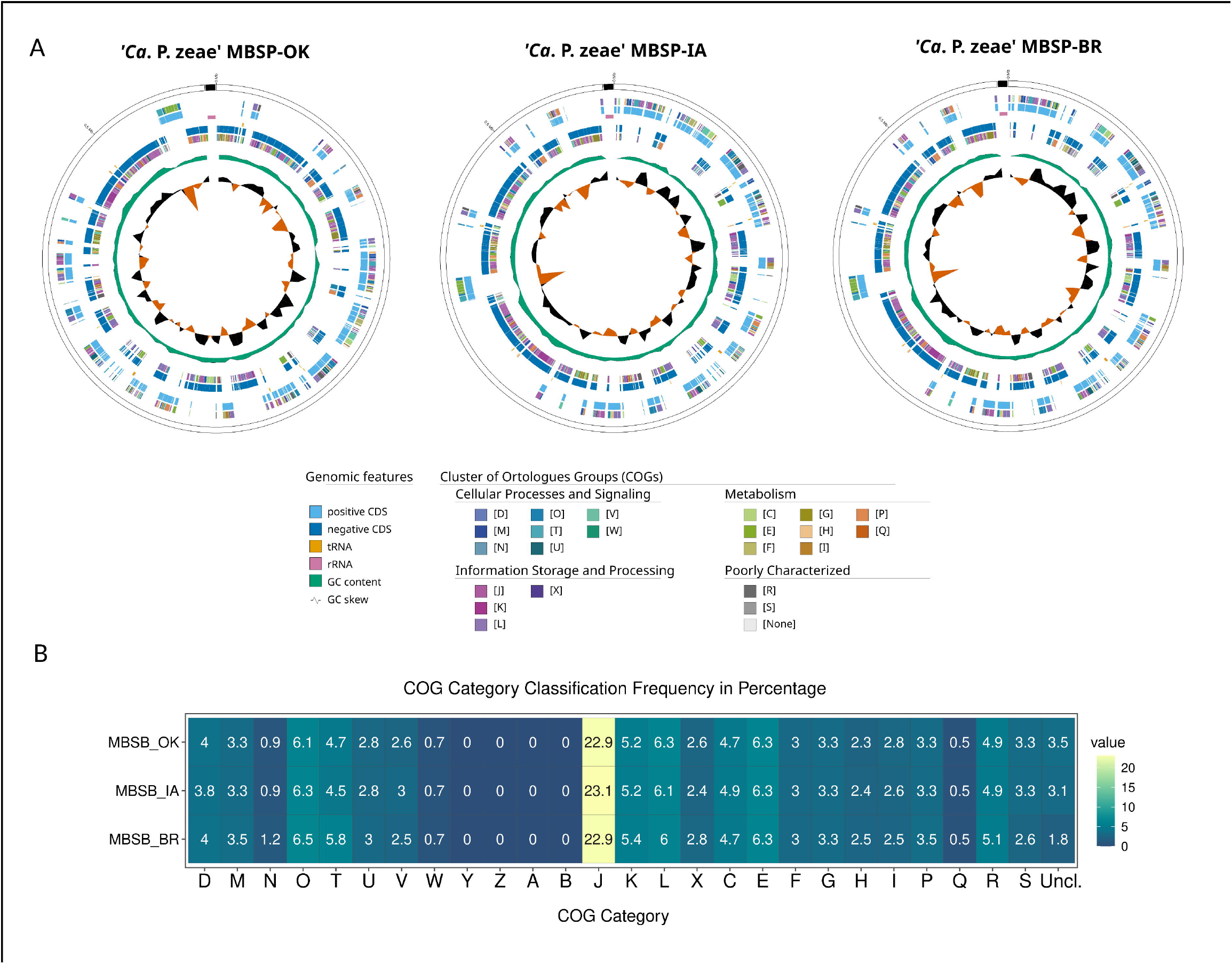
Maize bushy stunt phytoplasma genomes generated in this study. **A**) Circular genome maps of three *‘Ca. P. zeae’* strains, MBSP-OK, MBSP-IA, and MBSP-BR, showing coding sequences on the forward (blue) and reverse (light blue) strands, tRNA (orange), rRNA (red), GC content (black), and GC skew (green and orange). Outer rings display functional annotation by COG categories. **B**) Heatmap showing the frequency distribution of genes assigned to COG functional categories for each genome. Most genes are involved in information storage and processing, particularly translation (J), followed by genes related to replication (L) and transcription (K), consistent with genome reduction and adaptation to obligate host association.

To further resolve the phylogenetic position of MBSP strains, we constructed two phylogenies. The first was a phylogenomic tree based on concatenated single-copy orthologous genes from all available ‘*Ca*. Phytoplasma’ recognized species with genomes available in NCBI using GToTree [58]. The second tree was based on full-length 16S rRNA operon sequences aligned with MAFFT [59] and analyzed using IQ-TREE2 [60]. In both trees, all MBSP strains clustered in a well-supported monophyletic group with ‘*Ca*. Phytoplasma asteris’, clearly distinct from other closely related species such as ‘*Ca*. P. pruni’ and ‘*Ca*. P. mali’ (**Figure 3**). This grouping was supported by strong bootstrap values, reinforcing the phylogenetic placement despite the relatively small sequence divergence. This topology supports their classification within the 16SrI group while highlighting genomic divergence from other members. The 16S rRNA-based phylogeny, which included strains MBS-M3 and two newly reported genomes alongside 57 described ‘*Ca*. Phytoplasma’ species, showed that MBSP strains are clearly distinct from ‘*Ca*. P. asteris’, despite close sequence similarity (**Figure 4**). A common ancestor was evident for ‘*Ca*. P. zeae’, ‘*Ca*. P. asteris’, ‘*Ca*. P. triticii’, and ‘*Ca*. P. lycopersici’. Notably, ‘*Ca*. P. lycopersici’, like ‘*Ca*. P. zeae’, has only been reported in the Americas.

**Figure 3.**
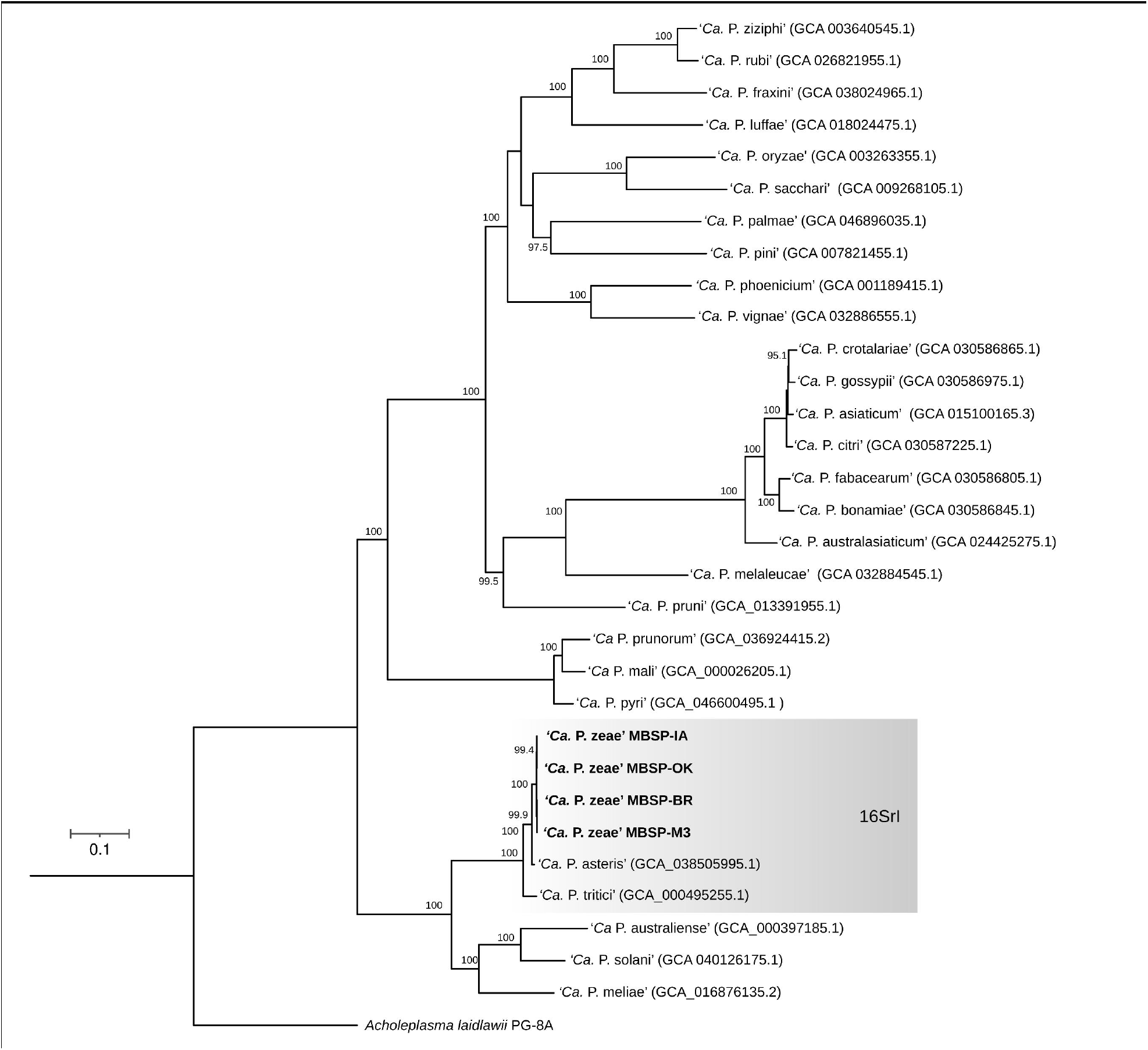
Phylogenomic tree based on concatenated single-copy orthologous genes from all available *‘Candidatus* Phytoplasma*’* genomes. The tree was inferred using maximum likelihood with bootstrap values shown at each node. Strains of *‘Ca. P. zeae’* (MBSP-IA, MBSP-OK, MBSP-BR, and MBSP-M3) clustered within a well-supported monophyletic clade alongside *‘Ca. P. asteris’* and *‘Ca. P. tritici’*, forming a subgroup within the 16SrI lineage. The tree was rooted with *Acholeplasma laidlawii* PG-8A as outgroup. Bootstrap support values ≥ 95% indicate high confidence in branching patterns.

**Figure 4.**
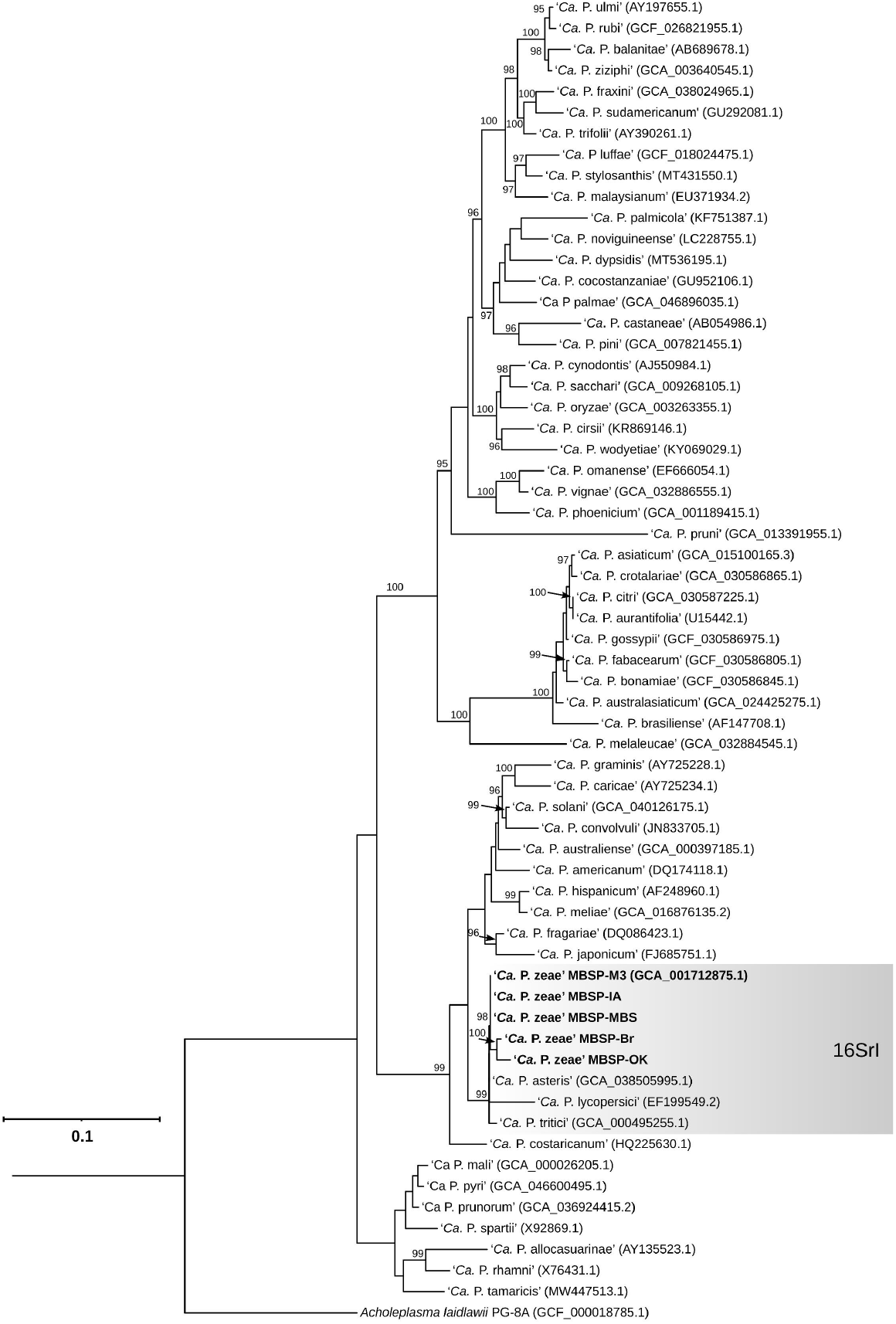
Phylogenetic tree based on full-length 16S rRNA gene sequences from *‘Candidatus* Phytoplasma*’* species, including the newly sequenced *‘Ca. P. zeae’* strains (MBSP-M3, MBSP-IA, MBSP-BR, and MBSP-OK). The tree was constructed using maximum likelihood and rooted with *Acholeplasma laidlawii* PG-8A as outgroup. The 16SrI clade, to which *‘Ca. P. zeae’* and *‘Ca. P. asteris’* belong, is shaded in gray. Bootstrap values (>95%) are shown at the nodes, indicating the robustness of each branch. This analysis confirms the close relationship between *‘Ca. P. zeae’* and other members of the 16SrI group, while also supporting its divergence from other phytoplasma species.

Pairwise identity analysis of 16S rRNA genes revealed > 99.8% identity between MBSP and ‘*Ca*. P. asteris’, but <97% identity with other formally described ‘*Ca*. Phytoplasma’ species. While MBSP and ‘*Ca*. P. asteris’ share specific diagnostic regions within the 16S rRNA gene [10], distinct nucleotide patterns were observed in ‘*Ca*. P. zeae’-related strains (**Table 1**). Eleven polymorphic sites differentiated ‘*Ca*. P. zeae’ from ‘*Ca*. P. asteris’, notably at positions 1319–1321, 1367–1369, and 151– 153 (**Table 1**). The 16S rRNA gene showed >99% identity between ‘*Ca*. P. zeae’ and ‘*Ca*. P. asteris’, but lower identity with ‘*Ca*. P. triticii’ (98.82%) and ‘*Ca*. P. lycopersici’ (96.71%). Given the subtle differences and high 16S similarity (>97.5%), we examined additional conserved genes for improved resolution.

**Table 1.** Nucleotide identity and unique sequences in MBS-Brasil^RS^ phytoplasma, reference strain of ‘*Candidatus* Phytoplasma zeae’, in comparison with ‘*Ca*. P. asteris’, ‘*Ca*. P. triticii’, and ‘*Ca*. P. lycopersici’.

Among the protein-coding genes analyzed, *rpoB* and *rps3* provided the best discriminatory power (**Table 1**). Nucleotide identity values for ‘*Ca*. P. zeae’ vs. ‘*Ca*. P. asteris’ were 99.31% (*rpoB*) and 99.34% (*rps3*), while identity dropped to 95.93% and 96.17%, respectively, when compared to ‘*Ca*. P. triticii’. The *rpoB* gene contained five distinct nucleotide differences (e.g., at positions 1268–1270 and 2850–2852), while *rps3* presented six (e.g., 17–19 and 570–572) (**Table 1**). Although *rpoB* is not part of the IRPCM-recognized minimal set for ‘*Ca*. Phytoplasma’ classification, it has been successfully used to distinguish closely related ‘*Ca*. P. asteris’ strains [42,61]. The analysis of the *cpn60*UT revealed three SNPs at positions 61, 394, and 1075, supporting its continued relevance in phylogenetic classification schemes, even though we did not detect the polymorphism at position 125 previously reported for MBS-Ver [62]. The *secY* and *secA* genes also demonstrated high identity (99.4% and 99.3%, respectively, vs. ‘*Ca*. P. asteris’) but showed lower similarity with ‘*Ca*. P. triticii’ (∼95.9%), each with nine informative differences.

Pangenome analysis reinforced these findings. Among 3,720 identified gene clusters (GCs), 276 were shared across all strains (core GCs), while 45 accessory GCs were exclusive to ‘*Ca*. P. zeae’. Among the accessory gene clusters, we identified genes associated with amino acid transport and metabolism (15 genes), defense mechanisms (8 genes), and posttranslational modification, protein turnover, and chaperones (2 genes). A substantial fraction (137 genes) lacked functional annotation **(Table S5)**. The pangenome clustering based on shared gene clusters content aligned with the phylogenetic patterns, with ‘*Ca*. P. zeae’ grouping alongside ‘*Ca*. P. asteris’ and ‘*Ca*. P. triticii’ **(Figure 5A)**. ANI analyses confirmed this relationship, showing 97.7–98% identity between ‘*Ca*. P. zeae’ and ‘*Ca*. P. asteris’, and 93.3–93.4% between ‘*Ca*. P. zeae’ and ‘*Ca*. P. triticii’ **(Figure 5B-C, Table S6)**. AAI analyses revealed a similar pattern: ‘*Ca*. P. zeae’ and ‘*Ca*. P. asteris’ shared high similarity, with AAI values ranging from 96.65% to 96.88%, whereas ‘*Ca*. P. triticii’ exhibited lower values, between 91.62% and 91.90% **(Table S7)**.

**Figure 5.**
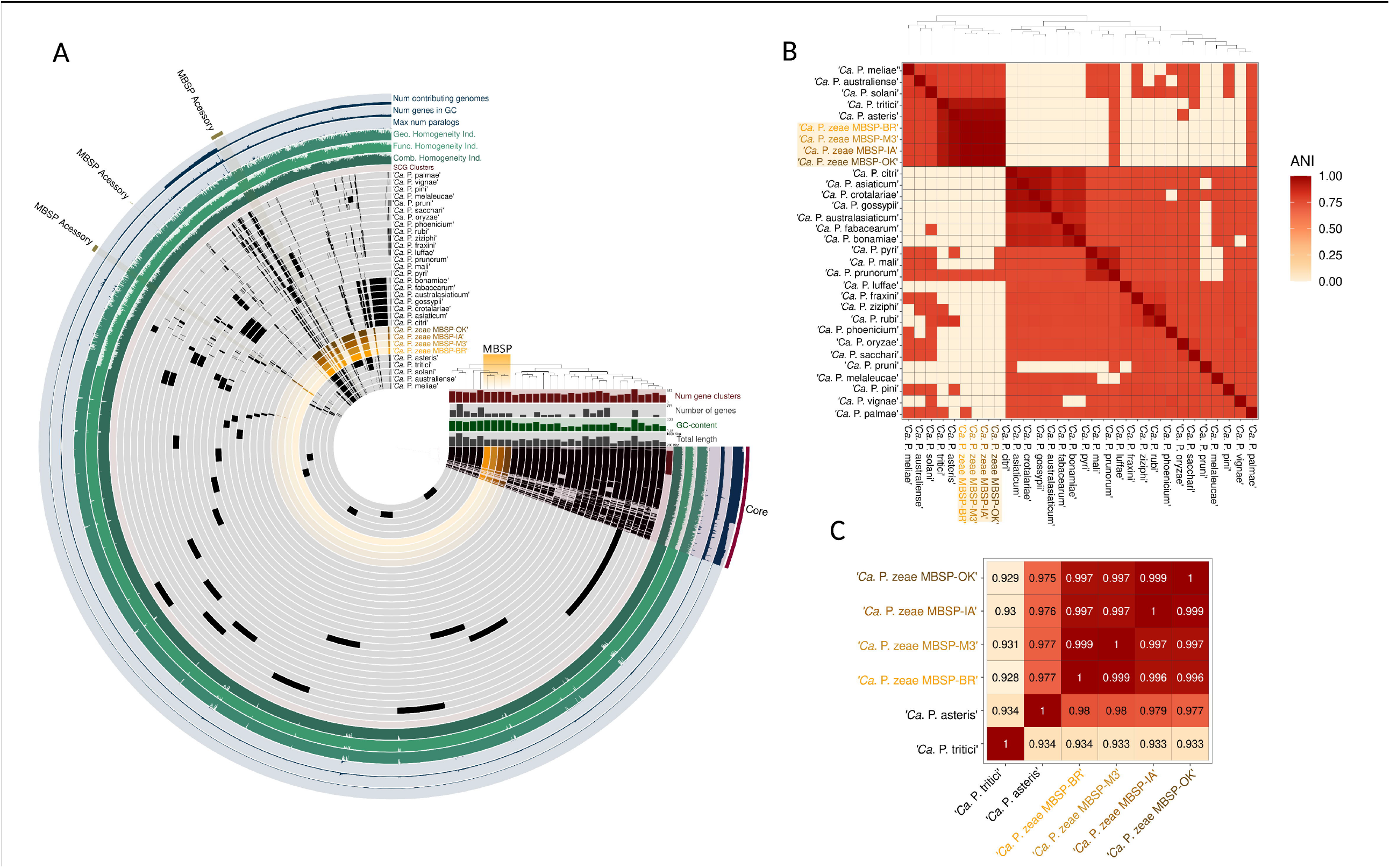
Comparative genomic analysis of *‘Ca*. P. *zeae’* and closely related *‘Candidatus* Phytoplasma*’* species. **A**) Pangenome clustering of 41 phytoplasma genomes based on presence/absence of gene clusters. Colored bars denote core and accessory gene sets, with MBSP-specific gene clusters highlighted in orange. Tracks display gene cluster metrics such as number of contributing genomes, number of genes, and functional homogeneity indices. **B**) Heatmap of Average Nucleotide Identity (ANI) values among the same genomes, showing high similarity among *‘Ca. P. zeae’* strains and clear separation from other taxa. **C**) ANI pairwise comparisons highlighting the high genomic similarity (>99.7%) among *‘Ca. P. zeae’* strains and their distinction from *‘Ca. P. asteris’* and *‘Ca. P. tritici’*, which share ANI values around 93.3–98% with MBSP genomes.

Despite the relatively limited sequence divergence among these taxa, the distinct ecological traits of ‘*Candidatus* Phytoplasma zeae’, including its specific insect vector, narrow host range, and geographically restricted distribution, provide strong support for its recognition as a separate species. This interpretation aligns with the ecological species criterium, which is particularly useful when genomic metrics such as ANI and AAI approach the thresholds for species delimitation. In this context, the consistent biological, geographic, and epidemiological distinctions justify its delineation as a new taxon. The authors proposing this classification represent a broad coalition of researchers who have studied maize bushy stunt phytoplasma for over a decade in Brazil, Colombia, Mexico, and the United States, together with a new generation of scientists working directly with growers, industry, stakeholders, and policymakers. This collective expertise and engagement lend strong support to the formal recognition of ‘*Ca*. P. zeae’.

### DESCRIPTION OF ‘CANDIDATUS PHYTOPLASMA ZEAE’

Based on the unique host (*Z. mays*), unique vectors (*D. maidis* and *D. elimatus)*, the geographic distribution restricted to the Americas, differentiating MBS phytoplasma from other ‘*Candidatus* Phytoplasma asteris’-related strains ecologically, and the consensus from the community represented by the authors, we propose the designation of the novel ‘*Candidatus* Phytoplasma zeae’ species described as detailed below:

‘*Candidatus* Phytoplasma zeae’ *(ze’ae*. L. gen. n. *zeae*, of spelt, of *Zea mays*, the known plant host of the phytoplasma).

### MBSP-BR^RS^ is the reference strain

[(Mollicutes) NC; NA; O, wall-less; NAS (Genome GenBank accession xxx); unique regions in the 16S rRNA-encoding gene are: 152, 337,359,370,507,532,537,1320,1368,1373,1425. Unique sequences *rpoB, rps19, rps3, rpl22, secY, secA, cpn60* or *GroEL* differences are shown in **Table 1**.

## SUPPLEMENTARY FILES

**Table S1**. Occurrences of *Dalbulus maidis, D. elimatus*, and Maize bushy stunt (MBS) phytoplasma. When a reference only provided the locality name (without coordinates), the record was registered based on this locality, not the exact point. For the MBS phytoplasma, only references that included the molecular identification level were included. Those based solely on symptomatic plants were excluded due to similarities with corn stunt spiroplasma symptoms.

**Table S2**. Detailed information on the samples analyzed in this study for the detection of ‘*Candidatus* Phytoplasma zeae’ and genome assemblies.

**Table S3**. Assembly statistics and genome quality metrics for *‘Candidatus* Phytoplasma*’* strains MBSP-BR, MBSP-IA, and MBSP-OK.

**Table S4**. Distribution of predicted protein-coding genes in *‘Candidatus* Phytoplasma*’* strains MBSP-BR, MBSP-IA, and MBSP-OK across functional categories defined by the Clusters of Orthologous Groups (COG) classification. Categories are grouped into Cellular Processes and Signaling, Information Storage and Processing, Metabolism, and Poorly Characterized, with the corresponding COG code and number of genes assigned to each category.

**Table S5**. Core and accessory gene clusters of *‘Candidatus* Phytoplasma zeae*’* strains MBSP-BR, MBSP-IA, and MBSP-OK, in comparison with gene clusters from other *‘Candidatus* Phytoplasma*’* species.

**Table S6**. Pairwise average amino acid identity (AAI) comparisons between *‘Candidatus* Phytoplasma zeae*’* strains MBSP-BR, MBSP-IA, and MBSP-OK and other *‘Ca*. Phytoplasma*’* genomes.

**Table S7**. Matrix of average nucleotide identity (ANI) values among *‘Candidatus* Phytoplasma zeae*’* strains MBSP-BR, MBSP-IA, and MBSP-OK, and representative genomes from other *‘Candidatus* Phytoplasma*’* species.

