## Supplementary material for "‘*Candidatus* Phytoplasma zeae’: community-driven species delineation of the maize bushy stunt phytoplasma, a *Dalbulus*-transmitted corn pathogen confined to the Americas": Table S1

**Table S1**. Occurrences of *Dalbulus maidis, D. elimatus,* and Maize bushy stunt (MBS) phytoplasma. When a reference only provided the locality name (without coordinates), the record was registered based on this locality, not the exact point. For the MBS phytoplasma, only references that included the molecular identification level were included. Those based solely on symptomatic plants were excluded due to similarities with corn stunt spiroplasma symptoms.

| **Species** | **Latitude** | **Longitude** | **Country** | **Reference** |
| --- | --- | --- | --- | --- |
| *D. maidis* | -32.60432 | -64.38400 | Argentina | Virla et al. 2010; Virla et al. 2013 |
| *D. maidis* | -31.06889 | -64.12670 | Argentina | Virla et al. 2010; Virla et al. 2013 |
| *D. maidis* | -30.92053 | -64.07542 | Argentina | Virla et al. 2010; Virla et al. 2013 |
| *D. maidis* | -29.89343 | -63.72887 | Argentina | Virla et al. 2010; Virla et al. 2013 |
| *D. maidis* | -27.07337 | -61.07945 | Argentina | Virla et al. 2010; Virla et al. 2013 |
| *D. maidis* | -26.83430 | -65.27381 | Argentina | Virla et al. 2010; Virla et al. 2013 |
| *D. maidis* | -26.60750 | -64.80910 | Argentina | Santana Jr et al. 2019 |
| *D. maidis* | -26.65040 | -64.54690 | Argentina | Santana Jr et al. 2019 |
| *D. maidis* | -26.55070 | -64.64080 | Argentina | Santana Jr et al. 2019 |
| *D. maidis* | -26.45410 | -64.56370 | Argentina | Santana Jr et al. 2019 |
| *D. maidis* | -26.35090 | -64.63030 | Argentina | Santana Jr et al. 2019 |
| *D. maidis* | -26.24900 | -64.52470 | Argentina | Santana Jr et al. 2019 |
| *D. maidis* | -26.02220 | -64.22620 | Argentina | Santana Jr et al. 2019 |
| *D. maidis* | -25.96320 | -63.95830 | Argentina | Santana Jr et al. 2019 |
| *D. maidis* | -26.17850 | -63.85010 | Argentina | Santana Jr et al. 2019 |
| *D. maidis* | -25.64420 | -64.08810 | Argentina | Santana Jr et al. 2019 |
| *D. maidis* | -25.75620 | -64.55930 | Argentina | Santana Jr et al. 2019 |
| *D. maidis* | -25.78530 | -64.81340 | Argentina | Santana Jr et al. 2019 |
| *D. maidis* | -25.61170 | -64.67000 | Argentina | Santana Jr et al. 2019 |
| *D. maidis* | -25.42550 | -64.81180 | Argentina | Santana Jr et al. 2019 |
| *D. maidis* | -25.13230 | -64.22660 | Argentina | Santana Jr et al. 2019 |
| *D. maidis* | -25.18250 | -63.78050 | Argentina | Santana Jr et al. 2019 |
| *D. maidis* | -25.20220 | -63.26650 | Argentina | Santana Jr et al. 2019 |
| *D. maidis* | -24.77810 | -62.84880 | Argentina | Santana Jr et al. 2019 |
| *D. maidis* | -24.41500 | -64.01950 | Argentina | Santana Jr et al. 2019 |
| *D. maidis* | -24.02410 | -63.92300 | Argentina | Santana Jr et al. 2019 |
| *D. maidis* | -23.62030 | -64.09760 | Argentina | Santana Jr et al. 2019 |
| *D. maidis* | -23.14390 | -63.84420 | Argentina | Santana Jr et al. 2019 |
| *D. maidis* | -22.60420 | -63.44600 | Argentina | Santana Jr et al. 2019 |
| *D. maidis* | -24.53050 | -62.08960 | Argentina | Santana Jr et al. 2019 |
| *D. maidis* | -24.87270 | -61.54070 | Argentina | Santana Jr et al. 2019 |
| *D. maidis* | -25.42120 | -61.86010 | Argentina | Santana Jr et al. 2019 |
| *D. maidis* | -25.78730 | -61.55420 | Argentina | Santana Jr et al. 2019 |
| *D. maidis* | -26.13370 | -60.97560 | Argentina | Santana Jr et al. 2019 |
| *D. maidis* | -26.61870 | -61.31730 | Argentina | Santana Jr et al. 2019 |
| *D. maidis* | -18.22506 | -63.84817 | Bolivia | Zahniser 2007 |
| *D. maidis* | -17.41839 | -66.20111 | Bolivia | Zahniser 2007 |
| *D. maidis* | -17.41667 | -63.25000 | Bolivia | Zahniser 2007 |
| *D. maidis* | -17.36347 | -66.32128 | Bolivia | Zahniser 2007 |
| *D. maidis* | -28.49078 | -52.55178 | Brazil | de Oliveira et al. 2007; Meneses et al. 2016; Oliveira et al. 2013a; Oliveira et al. 2013b; Waquil et al. 1999 |
| *D. maidis* | -28.13898 | -52.29003 | Brazil | de Oliveira et al. 2007; Meneses et al. 2016; Oliveira et al. 2013a; Oliveira et al. 2013b; Waquil et al. 1999 |
| *D. maidis* | -27.64573 | -53.32819 | Brazil | de Oliveira et al. 2007; Meneses et al. 2016; Oliveira et al. 2013a; Oliveira et al. 2013b; Waquil et al. 1999 |
| *D. maidis* | -27.37891 | -53.40049 | Brazil | de Oliveira et al. 2007; Meneses et al. 2016; Oliveira et al. 2013a; Oliveira et al. 2013b; Waquil et al. 1999 |
| *D. maidis* | -27.07296 | -52.70312 | Brazil | de Oliveira et al. 2007; Meneses et al. 2016; Oliveira et al. 2013a; Oliveira et al. 2013b; Waquil et al. 1999 |
| *D. maidis* | -27.04991 | -52.61833 | Brazil | de Oliveira et al. 2007; Meneses et al. 2016; Oliveira et al. 2013a; Oliveira et al. 2013b; Waquil et al. 1999 |
| *D. maidis* | -26.99751 | -51.87348 | Brazil | de Oliveira et al. 2007; Meneses et al. 2016; Oliveira et al. 2013a; Oliveira et al. 2013b; Waquil et al. 1999 |
| *D. maidis* | -26.89422 | -52.92124 | Brazil | de Oliveira et al. 2007; Meneses et al. 2016; Oliveira et al. 2013a; Oliveira et al. 2013b; Waquil et al. 1999 |
| *D. maidis* | -26.87883 | -53.17009 | Brazil | de Oliveira et al. 2007; Meneses et al. 2016; Oliveira et al. 2013a; Oliveira et al. 2013b; Waquil et al. 1999 |
| *D. maidis* | -26.82675 | -53.26586 | Brazil | de Oliveira et al. 2007; Meneses et al. 2016; Oliveira et al. 2013a; Oliveira et al. 2013b; Waquil et al. 1999 |
| *D. maidis* | -26.69481 | -53.51712 | Brazil | de Oliveira et al. 2007; Meneses et al. 2016; Oliveira et al. 2013a; Oliveira et al. 2013b; Waquil et al. 1999 |
| *D. maidis* | -26.58171 | -53.52164 | Brazil | de Oliveira et al. 2007; Meneses et al. 2016; Oliveira et al. 2013a; Oliveira et al. 2013b; Waquil et al. 1999 |
| *D. maidis* | -26.46770 | -53.50355 | Brazil | de Oliveira et al. 2007; Meneses et al. 2016; Oliveira et al. 2013a; Oliveira et al. 2013b; Waquil et al. 1999 |
| *D. maidis* | -26.39129 | -53.51970 | Brazil | de Oliveira et al. 2007; Meneses et al. 2016; Oliveira et al. 2013a; Oliveira et al. 2013b; Waquil et al. 1999 |
| *D. maidis* | -26.20907 | -53.63643 | Brazil | de Oliveira et al. 2007; Meneses et al. 2016; Oliveira et al. 2013a; Oliveira et al. 2013b; Waquil et al. 1999 |
| *D. maidis* | -26.07725 | -53.71153 | Brazil | de Oliveira et al. 2007; Meneses et al. 2016; Oliveira et al. 2013a; Oliveira et al. 2013b; Waquil et al. 1999 |
| *D. maidis* | -25.90569 | -50.44474 | Brazil | de Oliveira et al. 2007; Meneses et al. 2016; Oliveira et al. 2013a; Oliveira et al. 2013b; Waquil et al. 1999 |
| *D. maidis* | -25.75797 | -49.74333 | Brazil | de Oliveira et al. 2007; Meneses et al. 2016; Oliveira et al. 2013a; Oliveira et al. 2013b; Waquil et al. 1999 |
| *D. maidis* | -25.69775 | -53.77579 | Brazil | de Oliveira et al. 2007; Meneses et al. 2016; Oliveira et al. 2013a; Oliveira et al. 2013b; Waquil et al. 1999 |
| *D. maidis* | -25.02648 | -53.57596 | Brazil | de Oliveira et al. 2007; Meneses et al. 2016; Oliveira et al. 2013a; Oliveira et al. 2013b; Waquil et al. 1999 |
| *D. maidis* | -24.65008 | -53.74867 | Brazil | de Oliveira et al. 2007; Meneses et al. 2016; Oliveira et al. 2013a; Oliveira et al. 2013b; Waquil et al. 1999 |
| *D. maidis* | -24.64628 | -53.29883 | Brazil | de Oliveira et al. 2007; Meneses et al. 2016; Oliveira et al. 2013a; Oliveira et al. 2013b; Waquil et al. 1999 |
| *D. maidis* | -24.41319 | -52.81781 | Brazil | de Oliveira et al. 2007; Meneses et al. 2016; Oliveira et al. 2013a; Oliveira et al. 2013b; Waquil et al. 1999 |
| *D. maidis* | -23.99145 | -52.35483 | Brazil | de Oliveira et al. 2007; Meneses et al. 2016; Oliveira et al. 2013a; Oliveira et al. 2013b; Waquil et al. 1999 |
| *D. maidis* | -23.56396 | -51.53198 | Brazil | de Oliveira et al. 2007; Meneses et al. 2016; Oliveira et al. 2013a; Oliveira et al. 2013b; Waquil et al. 1999 |
| *D. maidis* | -23.51305 | -51.71274 | Brazil | de Oliveira et al. 2007; Meneses et al. 2016; Oliveira et al. 2013a; Oliveira et al. 2013b; Waquil et al. 1999 |
| *D. maidis* | -23.35021 | -51.89945 | Brazil | de Oliveira et al. 2007; Meneses et al. 2016; Oliveira et al. 2013a; Oliveira et al. 2013b; Waquil et al. 1999 |
| *D. maidis* | -23.25878 | -51.26167 | Brazil | de Oliveira et al. 2007; Meneses et al. 2016; Oliveira et al. 2013a; Oliveira et al. 2013b; Waquil et al. 1999 |
| *D. maidis* | -23.16638 | -50.61167 | Brazil | de Oliveira et al. 2007; Meneses et al. 2016; Oliveira et al. 2013a; Oliveira et al. 2013b; Waquil et al. 1999 |
| *D. maidis* | -23.14984 | -50.53593 | Brazil | de Oliveira et al. 2007; Meneses et al. 2016; Oliveira et al. 2013a; Oliveira et al. 2013b; Waquil et al. 1999 |
| *D. maidis* | -22.80343 | -47.62894 | Brazil | de Oliveira et al. 2007; Meneses et al. 2016; Oliveira et al. 2013a; Oliveira et al. 2013b; Waquil et al. 1999 |
| *D. maidis* | -22.25437 | -45.88173 | Brazil | de Oliveira et al. 2007; Meneses et al. 2016; Oliveira et al. 2013a; Oliveira et al. 2013b; Waquil et al. 1999 |
| *D. maidis* | -22.13533 | -45.74988 | Brazil | de Oliveira et al. 2007; Meneses et al. 2016; Oliveira et al. 2013a; Oliveira et al. 2013b; Waquil et al. 1999 |
| *D. maidis* | -21.90188 | -45.62874 | Brazil | de Oliveira et al. 2007; Meneses et al. 2016; Oliveira et al. 2013a; Oliveira et al. 2013b; Waquil et al. 1999 |
| *D. maidis* | -21.65465 | -45.31596 | Brazil | de Oliveira et al. 2007; Meneses et al. 2016; Oliveira et al. 2013a; Oliveira et al. 2013b; Waquil et al. 1999 |
| *D. maidis* | -21.40371 | -44.92868 | Brazil | de Oliveira et al. 2007; Meneses et al. 2016; Oliveira et al. 2013a; Oliveira et al. 2013b; Waquil et al. 1999 |
| *D. maidis* | -21.21078 | -45.05888 | Brazil | de Oliveira et al. 2007; Meneses et al. 2016; Oliveira et al. 2013a; Oliveira et al. 2013b; Waquil et al. 1999 |
| *D. maidis* | -21.01423 | -47.33700 | Brazil | de Oliveira et al. 2007; Meneses et al. 2016; Oliveira et al. 2013a; Oliveira et al. 2013b; Waquil et al. 1999 |
| *D. maidis* | -20.87736 | -46.37156 | Brazil | de Oliveira et al. 2007; Meneses et al. 2016; Oliveira et al. 2013a; Oliveira et al. 2013b; Waquil et al. 1999 |
| *D. maidis* | -20.50531 | -55.83438 | Brazil | de Oliveira et al. 2007; Meneses et al. 2016; Oliveira et al. 2013a; Oliveira et al. 2013b; Waquil et al. 1999 |
| *D. maidis* | -20.46559 | -45.80249 | Brazil | de Oliveira et al. 2007; Meneses et al. 2016; Oliveira et al. 2013a; Oliveira et al. 2013b; Waquil et al. 1999 |
| *D. maidis* | -19.86448 | -47.47283 | Brazil | de Oliveira et al. 2007; Meneses et al. 2016; Oliveira et al. 2013a; Oliveira et al. 2013b; Waquil et al. 1999 |
| *D. maidis* | -19.79104 | -45.63820 | Brazil | de Oliveira et al. 2007; Meneses et al. 2016; Oliveira et al. 2013a; Oliveira et al. 2013b; Waquil et al. 1999 |
| *D. maidis* | -19.57538 | -47.00335 | Brazil | de Oliveira et al. 2007; Meneses et al. 2016; Oliveira et al. 2013a; Oliveira et al. 2013b; Waquil et al. 1999 |
| *D. maidis* | -19.45972 | -44.17529 | Brazil | de Oliveira et al. 2007; Meneses et al. 2016; Oliveira et al. 2013a; Oliveira et al. 2013b; Waquil et al. 1999 |
| *D. maidis* | -19.18809 | -48.15771 | Brazil | de Oliveira et al. 2007; Meneses et al. 2016; Oliveira et al. 2013a; Oliveira et al. 2013b; Waquil et al. 1999 |
| *D. maidis* | -19.17739 | -47.68901 | Brazil | de Oliveira et al. 2007; Meneses et al. 2016; Oliveira et al. 2013a; Oliveira et al. 2013b; Waquil et al. 1999 |
| *D. maidis* | -18.11894 | -49.24617 | Brazil | de Oliveira et al. 2007; Meneses et al. 2016; Oliveira et al. 2013a; Oliveira et al. 2013b; Waquil et al. 1999 |
| *D. maidis* | -15.69038 | -47.78185 | Brazil | de Oliveira et al. 2007; Meneses et al. 2016; Oliveira et al. 2013a; Oliveira et al. 2013b; Waquil et al. 1999 |
| *D. maidis* | -8.49614 | -45.77746 | Brazil | de Oliveira et al. 2007; Meneses et al. 2016; Oliveira et al. 2013a; Oliveira et al. 2013b; Waquil et al. 1999 |
| *D. maidis* | -7.64006 | -46.34842 | Brazil | de Oliveira et al. 2007; Meneses et al. 2016; Oliveira et al. 2013a; Oliveira et al. 2013b; Waquil et al. 1999 |
| *D. maidis* | -7.57294 | -45.96202 | Brazil | de Oliveira et al. 2007; Meneses et al. 2016; Oliveira et al. 2013a; Oliveira et al. 2013b; Waquil et al. 1999 |
| *D. maidis* | -6.98939 | -45.43294 | Brazil | de Oliveira et al. 2007; Meneses et al. 2016; Oliveira et al. 2013a; Oliveira et al. 2013b; Waquil et al. 1999 |
| *D. maidis* | -5.23151 | -37.38243 | Brazil | de Oliveira et al. 2007; Meneses et al. 2016; Oliveira et al. 2013a; Oliveira et al. 2013b; Waquil et al. 1999 |
| *D. maidis* | -5.03159 | -42.79951 | Brazil | de Oliveira et al. 2007; Meneses et al. 2016; Oliveira et al. 2013a; Oliveira et al. 2013b; Waquil et al. 1999 |
| *D. maidis* | -17.87615 | -51.66616 | Brazil | de Oliveira et al. 2007; Meneses et al. 2016; Oliveira et al. 2013a; Oliveira et al. 2013b; Waquil et al. 1999 |
| *D. maidis* | -13.05170 | -55.87118 | Brazil | de Oliveira et al. 2007; Meneses et al. 2016; Oliveira et al. 2013a; Oliveira et al. 2013b; Waquil et al. 1999 |
| *D. maidis* | -16.80813 | -47.61953 | Brazil | de Oliveira et al. 2007; Meneses et al. 2016; Oliveira et al. 2013a; Oliveira et al. 2013b; Waquil et al. 1999 |
| *D. maidis* | -12.53862 | -55.67438 | Brazil | de Oliveira et al. 2007; Meneses et al. 2016; Oliveira et al. 2013a; Oliveira et al. 2013b; Waquil et al. 1999 |
| *D. maidis* | -19.78650 | -47.88372 | Brazil | de Oliveira et al. 2007; Meneses et al. 2016; Oliveira et al. 2013a; Oliveira et al. 2013b; Waquil et al. 1999 |
| *D. maidis* | -18.38343 | -52.66614 | Brazil | de Oliveira et al. 2007; Meneses et al. 2016; Oliveira et al. 2013a; Oliveira et al. 2013b; Waquil et al. 1999 |
| *D. maidis* | -17.80668 | -50.88779 | Brazil | de Oliveira et al. 2007; Meneses et al. 2016; Oliveira et al. 2013a; Oliveira et al. 2013b; Waquil et al. 1999 |
| *D. maidis* | -16.33276 | -46.91046 | Brazil | de Oliveira et al. 2007; Meneses et al. 2016; Oliveira et al. 2013a; Oliveira et al. 2013b; Waquil et al. 1999 |
| *D. maidis* | -12.39517 | -45.21213 | Brazil | de Oliveira et al. 2007; Meneses et al. 2016; Oliveira et al. 2013a; Oliveira et al. 2013b; Waquil et al. 1999 |
| *D. maidis* | -24.78629 | -49.77244 | Brazil | de Oliveira et al. 2007; Meneses et al. 2016; Oliveira et al. 2013a; Oliveira et al. 2013b; Waquil et al. 1999 |
| *D. maidis* | -25.41567 | -51.52491 | Brazil | de Oliveira et al. 2007; Meneses et al. 2016; Oliveira et al. 2013a; Oliveira et al. 2013b; Waquil et al. 1999 |
| *D. maidis* | -25.48727 | -50.62241 | Brazil | de Oliveira et al. 2007; Meneses et al. 2016; Oliveira et al. 2013a; Oliveira et al. 2013b; Waquil et al. 1999 |
| *D. maidis* | -23.98934 | -48.84611 | Brazil | de Oliveira et al. 2007; Meneses et al. 2016; Oliveira et al. 2013a; Oliveira et al. 2013b; Waquil et al. 1999 |
| *D. maidis* | -24.50791 | -50.39158 | Brazil | de Oliveira et al. 2007; Meneses et al. 2016; Oliveira et al. 2013a; Oliveira et al. 2013b; Waquil et al. 1999 |
| *D. maidis* | -17.29163 | -51.23430 | Brazil | de Oliveira et al. 2007; Meneses et al. 2016; Oliveira et al. 2013a; Oliveira et al. 2013b; Waquil et al. 1999 |
| *D. maidis* | -27.39065 | -51.24018 | Brazil | de Oliveira et al. 2007; Meneses et al. 2016; Oliveira et al. 2013a; Oliveira et al. 2013b; Waquil et al. 1999 |
| *D. maidis* | -24.88558 | -53.48617 | Brazil | de Oliveira et al. 2007; Meneses et al. 2016; Oliveira et al. 2013a; Oliveira et al. 2013b; Waquil et al. 1999 |
| *D. maidis* | -24.76918 | -51.73829 | Brazil | de Oliveira et al. 2007; Meneses et al. 2016; Oliveira et al. 2013a; Oliveira et al. 2013b; Waquil et al. 1999 |
| *D. maidis* | -12.58292 | -56.23058 | Brazil | de Oliveira et al. 2007; Meneses et al. 2016; Oliveira et al. 2013a; Oliveira et al. 2013b; Waquil et al. 1999 |
| *D. maidis* | -15.31531 | -54.78717 | Brazil | de Oliveira et al. 2007; Meneses et al. 2016; Oliveira et al. 2013a; Oliveira et al. 2013b; Waquil et al. 1999 |
| *D. maidis* | -23.83644 | -49.15704 | Brazil | de Oliveira et al. 2007; Meneses et al. 2016; Oliveira et al. 2013a; Oliveira et al. 2013b; Waquil et al. 1999 |
| *D. maidis* | -15.98570 | -47.56006 | Brazil | de Oliveira et al. 2007; Meneses et al. 2016; Oliveira et al. 2013a; Oliveira et al. 2013b; Waquil et al. 1999 |
| *D. maidis* | -13.56164 | -58.89434 | Brazil | de Oliveira et al. 2007; Meneses et al. 2016; Oliveira et al. 2013a; Oliveira et al. 2013b; Waquil et al. 1999 |
| *D. maidis* | -15.48625 | -54.29713 | Brazil | de Oliveira et al. 2007; Meneses et al. 2016; Oliveira et al. 2013a; Oliveira et al. 2013b; Waquil et al. 1999 |
| *D. maidis* | -22.27715 | -54.77588 | Brazil | de Oliveira et al. 2007; Meneses et al. 2016; Oliveira et al. 2013a; Oliveira et al. 2013b; Waquil et al. 1999 |
| *D. maidis* | -19.38624 | -47.35386 | Brazil | de Oliveira et al. 2007; Meneses et al. 2016; Oliveira et al. 2013a; Oliveira et al. 2013b; Waquil et al. 1999 |
| *D. maidis* | -25.12491 | -50.23779 | Brazil | de Oliveira et al. 2007; Meneses et al. 2016; Oliveira et al. 2013a; Oliveira et al. 2013b; Waquil et al. 1999 |
| *D. maidis* | -25.48198 | -51.90455 | Brazil | de Oliveira et al. 2007; Meneses et al. 2016; Oliveira et al. 2013a; Oliveira et al. 2013b; Waquil et al. 1999 |
| *D. maidis* | -18.73271 | -52.63871 | Brazil | de Oliveira et al. 2007; Meneses et al. 2016; Oliveira et al. 2013a; Oliveira et al. 2013b; Waquil et al. 1999 |
| *D. maidis* | -21.59274 | -55.16701 | Brazil | de Oliveira et al. 2007; Meneses et al. 2016; Oliveira et al. 2013a; Oliveira et al. 2013b; Waquil et al. 1999 |
| *D. maidis* | -16.29586 | -47.87334 | Brazil | de Oliveira et al. 2007; Meneses et al. 2016; Oliveira et al. 2013a; Oliveira et al. 2013b; Waquil et al. 1999 |
| *D. maidis* | -12.16623 | -45.72854 | Brazil | de Oliveira et al. 2007; Meneses et al. 2016; Oliveira et al. 2013a; Oliveira et al. 2013b; Waquil et al. 1999 |
| *D. maidis* | -17.50559 | -52.55606 | Brazil | de Oliveira et al. 2007; Meneses et al. 2016; Oliveira et al. 2013a; Oliveira et al. 2013b; Waquil et al. 1999 |
| *D. maidis* | -19.20680 | -54.59664 | Brazil | de Oliveira et al. 2007; Meneses et al. 2016; Oliveira et al. 2013a; Oliveira et al. 2013b; Waquil et al. 1999 |
| *D. maidis* | -17.51884 | -52.22939 | Brazil | de Oliveira et al. 2007; Meneses et al. 2016; Oliveira et al. 2013a; Oliveira et al. 2013b; Waquil et al. 1999 |
| *D. maidis* | -25.73546 | -51.70264 | Brazil | de Oliveira et al. 2007; Meneses et al. 2016; Oliveira et al. 2013a; Oliveira et al. 2013b; Waquil et al. 1999 |
| *D. maidis* | -22.62469 | -54.80386 | Brazil | de Oliveira et al. 2007; Meneses et al. 2016; Oliveira et al. 2013a; Oliveira et al. 2013b; Waquil et al. 1999 |
| *D. maidis* | -15.64023 | -46.45817 | Brazil | de Oliveira et al. 2007; Meneses et al. 2016; Oliveira et al. 2013a; Oliveira et al. 2013b; Waquil et al. 1999 |
| *D. maidis* | -18.54108 | -52.94955 | Brazil | de Oliveira et al. 2007; Meneses et al. 2016; Oliveira et al. 2013a; Oliveira et al. 2013b; Waquil et al. 1999 |
| *D. maidis* | -31.38726 | -52.71554 | Brazil | de Oliveira et al. 2007; Meneses et al. 2016; Oliveira et al. 2013a; Oliveira et al. 2013b; Waquil et al. 1999 |
| *D. maidis* | -26.15373 | -49.78481 | Brazil | de Oliveira et al. 2007; Meneses et al. 2016; Oliveira et al. 2013a; Oliveira et al. 2013b; Waquil et al. 1999 |
| *D. maidis* | -24.96368 | -50.12373 | Brazil | de Oliveira et al. 2007; Meneses et al. 2016; Oliveira et al. 2013a; Oliveira et al. 2013b; Waquil et al. 1999 |
| *D. maidis* | -25.35752 | -54.26924 | Brazil | de Oliveira et al. 2007; Meneses et al. 2016; Oliveira et al. 2013a; Oliveira et al. 2013b; Waquil et al. 1999 |
| *D. maidis* | -12.17932 | -45.00745 | Brazil | de Oliveira et al. 2007; Meneses et al. 2016; Oliveira et al. 2013a; Oliveira et al. 2013b; Waquil et al. 1999 |
| *D. maidis* | -21.76810 | -47.11003 | Brazil | de Oliveira et al. 2007; Meneses et al. 2016; Oliveira et al. 2013a; Oliveira et al. 2013b; Waquil et al. 1999 |
| *D. maidis* | -24.66452 | -50.85886 | Brazil | de Oliveira et al. 2007; Meneses et al. 2016; Oliveira et al. 2013a; Oliveira et al. 2013b; Waquil et al. 1999 |
| *D. maidis* | -18.40543 | -46.43889 | Brazil | de Oliveira et al. 2007; Meneses et al. 2016; Oliveira et al. 2013a; Oliveira et al. 2013b; Waquil et al. 1999 |
| *D. maidis* | -17.92380 | -50.84200 | Brazil | Santana Jr et al. 2019 |
| *D. maidis* | -17.84450 | -50.87100 | Brazil | Santana Jr et al. 2019 |
| *D. maidis* | -17.84370 | -50.93080 | Brazil | Santana Jr et al. 2019 |
| *D. maidis* | -17.86030 | -51.01700 | Brazil | Santana Jr et al. 2019 |
| *D. maidis* | -17.64210 | -51.10620 | Brazil | Santana Jr et al. 2019 |
| *D. maidis* | -17.63670 | -51.19470 | Brazil | Santana Jr et al. 2019 |
| *D. maidis* | -17.71120 | -51.20510 | Brazil | Santana Jr et al. 2019 |
| *D. maidis* | -17.79310 | -51.23140 | Brazil | Santana Jr et al. 2019 |
| *D. maidis* | -17.74170 | -51.33470 | Brazil | Santana Jr et al. 2019 |
| *D. maidis* | -17.65790 | -51.30010 | Brazil | Santana Jr et al. 2019 |
| *D. maidis* | -17.60650 | -51.39610 | Brazil | Santana Jr et al. 2019 |
| *D. maidis* | -17.45810 | -51.30050 | Brazil | Santana Jr et al. 2019 |
| *D. maidis* | -17.43310 | -51.50360 | Brazil | Santana Jr et al. 2019 |
| *D. maidis* | -17.43330 | -51.64580 | Brazil | Santana Jr et al. 2019 |
| *D. maidis* | -17.78280 | -51.56820 | Brazil | Santana Jr et al. 2019 |
| *D. maidis* | -17.72750 | -51.63690 | Brazil | Santana Jr et al. 2019 |
| *D. maidis* | -17.88870 | -51.82820 | Brazil | Santana Jr et al. 2019 |
| *D. maidis* | -17.95230 | -51.89770 | Brazil | Santana Jr et al. 2019 |
| *D. maidis* | -17.84550 | -51.94840 | Brazil | Santana Jr et al. 2019 |
| *D. maidis* | -17.83460 | -52.03110 | Brazil | Santana Jr et al. 2019 |
| *D. maidis* | -17.87330 | -52.06260 | Brazil | Santana Jr et al. 2019 |
| *D. maidis* | -17.94970 | -52.15570 | Brazil | Santana Jr et al. 2019 |
| *D. maidis* | -17.86310 | -52.16620 | Brazil | Santana Jr et al. 2019 |
| *D. maidis* | -17.74790 | -52.14970 | Brazil | Santana Jr et al. 2019 |
| *D. maidis* | -17.70420 | -52.21960 | Brazil | Santana Jr et al. 2019 |
| *D. maidis* | 3.51889 | -76.31594 | Colombia | Heady and Nault 1985; Triplehorn and Nault 1985 |
| *D. maidis* | 4.69154 | -74.20591 | Colombia | Heady and Nault 1985; Triplehorn and Nault 1985 |
| *D. maidis* | 4.71720 | -74.24725 | Colombia | Heady and Nault 1985; Triplehorn and Nault 1985 |
| *D. maidis* | 6.13049 | -75.35549 | Colombia | Heady and Nault 1985; Triplehorn and Nault 1985 |
| *D. maidis* | 6.30842 | -75.59115 | Colombia | Heady and Nault 1985; Triplehorn and Nault 1985 |
| *D. maidis* | 6.53875 | -75.83035 | Colombia | Heady and Nault 1985; Triplehorn and Nault 1985 |
| *D. maidis* | 6.95719 | -75.42146 | Colombia | Heady and Nault 1985; Triplehorn and Nault 1985 |
| *D. maidis* | 9.96615 | -84.21354 | Costa Rica | Heady and Nault 1985 |
| *D. maidis* | 10.00620 | -84.26441 | Costa Rica | Heady and Nault 1985 |
| *D. maidis* | 10.19181 | -84.38720 | Costa Rica | Heady and Nault 1985 |
| *D. maidis* | 10.49694 | -84.52276 | Costa Rica | Heady and Nault 1985 |
| *D. maidis* | 21.71020 | -82.76160 | Cuba | Genaro and Portuondo 1997 |
| *D. maidis* | 13.45011 | -89.12415 | El Salvador | Quezada 1979 |
| *D. maidis* | 14.17449 | -91.31297 | Guatemala | Zahniser 2007 |
| *D. maidis* | 14.30028 | -91.57344 | Guatemala | Zahniser 2007 |
| *D. maidis* | 14.33032 | -89.69997 | Guatemala | Zahniser 2007 |
| *D. maidis* | 14.33736 | -91.03621 | Guatemala | Zahniser 2007 |
| *D. maidis* | 14.06120 | -86.29678 | Honduras | Zahniser 2007 |
| *D. maidis* | 14.06345 | -86.86405 | Honduras | Zahniser 2007 |
| *D. maidis* | 14.13533 | -86.24937 | Honduras | Zahniser 2007 |
| *D. maidis* | 14.27982 | -87.40029 | Honduras | Zahniser 2007 |
| *D. maidis* | 17.99087 | -76.89595 | Jamaica | Eden‐Green and Waters 1981 |
| *D. maidis* | 17.00201 | -100.07512 | Mexico | Bellota et al. 2018; Hidalgo et al. 1998; Moya-Raygoza 2012 |
| *D. maidis* | 17.02384 | -97.93935 | Mexico | Bellota et al. 2018; Hidalgo et al. 1998; Moya-Raygoza 2012 |
| *D. maidis* | 17.97825 | -97.86692 | Mexico | Bellota et al. 2018; Hidalgo et al. 1998; Moya-Raygoza 2012 |
| *D. maidis* | 18.36241 | -99.87896 | Mexico | Bellota et al. 2018; Hidalgo et al. 1998; Moya-Raygoza 2012 |
| *D. maidis* | 18.64850 | -97.38848 | Mexico | Bellota et al. 2018; Hidalgo et al. 1998; Moya-Raygoza 2012 |
| *D. maidis* | 18.81701 | -97.20934 | Mexico | Bellota et al. 2018; Hidalgo et al. 1998; Moya-Raygoza 2012 |
| *D. maidis* | 18.83663 | -97.07482 | Mexico | Bellota et al. 2018; Hidalgo et al. 1998; Moya-Raygoza 2012 |
| *D. maidis* | 18.83969 | -96.41044 | Mexico | Bellota et al. 2018; Hidalgo et al. 1998; Moya-Raygoza 2012 |
| *D. maidis* | 19.01535 | -98.27882 | Mexico | Bellota et al. 2018; Hidalgo et al. 1998; Moya-Raygoza 2012 |
| *D. maidis* | 19.12471 | -96.14997 | Mexico | Bellota et al. 2018; Hidalgo et al. 1998; Moya-Raygoza 2012 |
| *D. maidis* | 19.53172 | -98.84592 | Mexico | Bellota et al. 2018; Hidalgo et al. 1998; Moya-Raygoza 2012 |
| *D. maidis* | 19.79331 | -104.21652 | Mexico | Bellota et al. 2018; Hidalgo et al. 1998; Moya-Raygoza 2012 |
| *D. maidis* | 19.87824 | -103.57315 | Mexico | Bellota et al. 2018; Hidalgo et al. 1998; Moya-Raygoza 2012 |
| *D. maidis* | 12.33750 | -86.68568 | Nicaragua | Power 1989 |
| *D. maidis* | -14.83840 | -74.95657 | Peru | Nault et al. 1981 |
| *D. maidis* | -14.08513 | -75.67685 | Peru | Nault et al. 1981 |
| *D. maidis* | -13.50392 | -71.98201 | Peru | Nault et al. 1981 |
| *D. maidis* | -13.42854 | -76.15044 | Peru | Nault et al. 1981 |
| *D. maidis* | -13.32306 | -72.12728 | Peru | Nault et al. 1981 |
| *D. maidis* | -13.31969 | -72.08228 | Peru | Nault et al. 1981 |
| *D. maidis* | -13.26032 | -72.26837 | Peru | Nault et al. 1981 |
| *D. maidis* | -13.16857 | -74.18673 | Peru | Nault et al. 1981 |
| *D. maidis* | -13.11426 | -74.19028 | Peru | Nault et al. 1981 |
| *D. maidis* | -11.99103 | -76.92950 | Peru | Nault et al. 1981 |
| *D. maidis* | -7.33324 | -78.17762 | Peru | Nault et al. 1981 |
| *D. maidis* | -7.31868 | -78.17204 | Peru | Nault et al. 1981 |
| *D. maidis* | -7.24388 | -78.38309 | Peru | Nault et al. 1981 |
| *D. maidis* | -7.19995 | -78.32386 | Peru | Nault et al. 1981 |
| *D. maidis* | -7.15629 | -78.49803 | Peru | Nault et al. 1981 |
| *D. maidis* | -7.07980 | -76.51103 | Peru | Nault et al. 1981 |
| *D. maidis* | 17.97287 | -66.26129 | Puerto Rico | Zahniser 2007 |
| *D. maidis* | 26.14252 | -98.22472 | United States | Summers et al. 2004 |
| *D. maidis* | 26.60454 | -80.58271 | United States | Summers et al. 2004 |
| *D. maidis* | 31.31276 | -92.51217 | United States | Summers et al. 2004 |
| *D. maidis* | 31.45392 | -83.46290 | United States | Summers et al. 2004 |
| *D. maidis* | 33.66018 | -93.66747 | United States | Summers et al. 2004 |
| *D. maidis* | 33.86385 | -91.44481 | United States | Summers et al. 2004 |
| *D. maidis* | 35.25073 | -92.69142 | United States | Summers et al. 2004 |
| *D. maidis* | 36.11005 | -119.89433 | United States | Summers et al. 2004 |
| *D. maidis* | 36.11837 | -118.86043 | United States | Summers et al. 2004 |
| *D. maidis* | 36.75994 | -119.51644 | United States | Summers et al. 2004 |
| *D. maidis* | 8.79502 | -69.75772 | Venezuela | Clavijo and Notz 1978 |
| *D. maidis* | 9.91937 | -67.38934 | Venezuela | Clavijo and Notz 1978 |
| *D. maidis* | 4.61979 | -76.01779 | Colombia | Arcila Bohórquez et al. 2024 |
| *D. maidis* | 4.61464 | -76.00470 | Colombia | Arcila Bohórquez et al. 2024 |
| *D. maidis* | 4.62045 | -75.99589 | Colombia | Arcila Bohórquez et al. 2024 |
| *D. maidis* | 4.58304 | -76.05802 | Colombia | Arcila Bohórquez et al. 2024 |
| *D. maidis* | 4.62244 | -76.02020 | Colombia | Arcila Bohórquez et al. 2024 |
| *D. maidis* | 4.62244 | -76.02020 | Colombia | Arcila Bohórquez et al. 2024 |
| *D. maidis* | 4.62244 | -76.02020 | Colombia | Arcila Bohórquez et al. 2024 |
| *D. maidis* | 3.69702 | -76.34142 | Colombia | Arcila Bohórquez et al. 2024 |
| *D. maidis* | 3.66410 | -73.57810 | Colombia | Arcila Bohórquez et al. 2024 |
| *D. maidis* | 8.93044 | -75.78902 | Colombia | Arcila Bohórquez et al. 2024 |
| *D. maidis* | 3.99985 | -73.38125 | Colombia | Arcila Bohórquez et al. 2024 |
| *D. maidis* | 4.18639 | -74.96396 | Colombia | Arcila Bohórquez et al. 2024 |
| *D. maidis* | 4.19809 | -74.90603 | Colombia | Arcila Bohórquez et al. 2024 |
| *D. maidis* | 3.66437 | -73.57928 | Colombia | Arcila Bohórquez et al. 2024 |
| *D. maidis* | 4.21408 | -72.48489 | Colombia | Arcila Bohórquez et al. 2024 |
| *D. maidis* | 8.92274 | -75.83205 | Colombia | Arcila Bohórquez et al. 2024 |
| *D. maidis* | 3.51227 | -76.28361 | Colombia | Arcila Bohórquez et al. 2024 |
| *D. maidis* | 4.08928 | -74.92521 | Colombia | Arcila Bohórquez et al. 2024 |
| *D. maidis* | 3.50769 | -73.72275 | Colombia | Arcila Bohórquez et al. 2024 |
| *D. maidis* | 3.51141 | -76.33840 | Colombia | Arcila Bohórquez et al. 2024 |
| *D. maidis* | 4.37619 | -76.14038 | Colombia | Arcila Bohórquez et al. 2024 |
| *D. maidis* | 8.93050 | -75.78886 | Colombia | Arcila Bohórquez et al. 2024 |
| *D. maidis* | 4.17862 | -74.89360 | Colombia | Arcila Bohórquez et al. 2024 |
| *D. maidis* | 4.45448 | -71.52541 | Colombia | Arcila Bohórquez et al. 2024 |
| *D. maidis* | 7.89149 | -73.50061 | Colombia | Arcila Bohórquez et al. 2024 |
| *D. maidis* | 10.63308 | -73.12482 | Colombia | Arcila Bohórquez et al. 2024 |
| *D. maidis* | 2.668382 | -75.333302 | Colombia | Reinoso, 2021 |
| *D. maidis* | 2.869621 | -75.281099 | Colombia | Reinoso, 2021 |
| *D. maidis* | 2.225862 | -75.611987 | Colombia | Reinoso, 2021 |
| *D. maidis* | 2.108519 | -75.831831 | Colombia | Reinoso, 2021 |
| *D. maidis* | 12.14722 | -86.16278 | Nicaragua | GBIF, 2025 |
| *D. maidis* | 43.08991 | -89.37417 | United States | GBIF, 2025 |
| *D. maidis* | 43.08991 | -89.37417 | United States | GBIF, 2025 |
| *D. maidis* | 43.30743 | -89.91241 | United States | GBIF, 2025 |
| *D. maidis* | 39.52448 | -79.81770 | United States | GBIF, 2025 |
| *D. maidis* | 39.23004 | -86.42012 | United States | GBIF, 2025 |
| *D. maidis* | 39.52443 | -79.81761 | United States | GBIF, 2025 |
| *D. maidis* | 42.61599 | -83.19203 | United States | GBIF, 2025 |
| *D. maidis* | 42.20844 | -93.42352 | United States | GBIF, 2025 |
| *D. maidis* | 41.21864 | -75.84174 | United States | GBIF, 2025 |
| *D. maidis* | 39.52441 | -79.81753 | United States | GBIF, 2025 |
| *D. maidis* | 39.52448 | -79.81770 | United States | GBIF, 2025 |
| *D. maidis* | 42.58745 | -83.19291 | United States | GBIF, 2025 |
| *D. maidis* | 39.52443 | -79.81757 | United States | GBIF, 2025 |
| *D. maidis* | 35.02559 | -85.36419 | United States | GBIF, 2025 |
| *D. maidis* | 35.14585 | -79.01185 | United States | GBIF, 2025 |
| *D. maidis* | -7.32944 | -71.84611 | Brazil | GBIF, 2025 |
| *D. maidis* | -31.38078 | -64.20966 | Argentina | GBIF, 2025 |
| *D. maidis* | 30.35750 | -97.72340 | United States | GBIF, 2025 |
| *D. maidis* | 32.24645 | -110.88481 | United States | GBIF, 2025 |
| *D. maidis* | -27.57670 | -56.67837 | Argentina | GBIF, 2025 |
| *D. maidis* | 37.50326 | -119.96137 | United States | GBIF, 2025 |
| *D. maidis* | 28.26229 | -105.49278 | México | GBIF, 2025 |
| *D. maidis* | 16.81667 | -92.96667 | México | GBIF, 2025 |
| *D. maidis* | 33.26590 | -88.87070 | United States | GBIF, 2025 |
| *D. maidis* | 33.26590 | -88.87070 | United States | GBIF, 2025 |
| *D. maidis* | 16.81667 | -92.96667 | México | GBIF, 2025 |
| *D. maidis* | 33.26590 | -88.87070 | United States | GBIF, 2025 |
| *D. maidis* | 33.26590 | -88.87070 | United States | GBIF, 2025 |
| *D. maidis* | 33.26590 | -88.87070 | United States | GBIF, 2025 |
| *D. maidis* | 33.26590 | -88.87070 | United States | GBIF, 2025 |
| *D. maidis* | 19.66833 | -104.03139 | México | GBIF, 2025 |
| *D. maidis* | 19.05000 | -104.26667 | México | GBIF, 2025 |
| *D. maidis* | 19.98333 | -97.18333 | México | GBIF, 2025 |
| *D. maidis* | 19.66833 | -104.03139 | México | GBIF, 2025 |
| *D. maidis* | 18.20130 | -67.14510 | Puerto Rico | GBIF, 2025 |
| *D. maidis* | 18.20130 | -67.14510 | Puerto Rico | GBIF, 2025 |
| *D. maidis* | 18.20130 | -67.14510 | Puerto Rico | GBIF, 2025 |
| *D. maidis* | 18.20130 | -67.14510 | Puerto Rico | GBIF, 2025 |
| *D. maidis* | 4.29860 | -74.80460 | Colombia | GBIF, 2025 |
| *D. maidis* | 18.20130 | -67.14510 | Puerto Rico | GBIF, 2025 |
| *D. maidis* | 19.48700 | -70.61500 | Dominican Republic | GBIF, 2025 |
| *D. maidis* | 18.20130 | -67.14510 | Puerto Rico | GBIF, 2025 |
| *D. maidis* | 18.20130 | -67.14510 | Puerto Rico | GBIF, 2025 |
| *D. maidis* | 19.48700 | -70.61500 | Dominican Republic | GBIF, 2025 |
| *D. maidis* | 4.29860 | -74.80460 | Colombia | GBIF, 2025 |
| *D. maidis* | 18.20130 | -67.14510 | Puerto Rico | GBIF, 2025 |
| *D. maidis* | 19.48700 | -70.61500 | Dominican Republic | GBIF, 2025 |
| *D. maidis* | 19.48700 | -70.61500 | Dominican Republic | GBIF, 2025 |
| *D. maidis* | 4.34500 | -74.36100 | Colombia | GBIF, 2025 |
| *D. maidis* | 19.48700 | -70.61500 | Dominican Republic | GBIF, 2025 |
| *D. maidis* | 19.48700 | -70.61500 | Dominican Republic | GBIF, 2025 |
| *D. maidis* | 18.20130 | -67.14510 | Puerto Rico | GBIF, 2025 |
| *D. maidis* | 19.48700 | -70.61500 | Dominican Republic | GBIF, 2025 |
| *D. maidis* | 15.43200 | -87.90400 | Honduras | GBIF, 2025 |
| *D. maidis* | 18.20130 | -67.14510 | Puerto Rico | GBIF, 2025 |
| *D. maidis* | 19.48700 | -70.61500 | Dominican Republic | GBIF, 2025 |
| *D. maidis* | 18.20130 | -67.14510 | Puerto Rico | GBIF, 2025 |
| *D. maidis* | 19.48700 | -70.61500 | Dominican Republic | GBIF, 2025 |
| *D. maidis* | 19.48700 | -70.61500 | Dominican Republic | GBIF, 2025 |
| *D. maidis* | 18.20130 | -67.14510 | Puerto Rico | GBIF, 2025 |
| *D. maidis* | 19.48700 | -70.61500 | Dominican Republic | GBIF, 2025 |
| *D. maidis* | 19.48700 | -70.61500 | Dominican Republic | GBIF, 2025 |
| *D. maidis* | 30.31070 | -91.09616 | United States | GBIF, 2025 |
| *D. maidis* | 40.35880 | -91.41880 | United States | GBIF, 2025 |
| *D. maidis* | 31.63464 | -98.24814 | United States | GBIF, 2025 |
| *D. maidis* | 31.63464 | -98.24814 | United States | GBIF, 2025 |
| *D. maidis* | 30.35750 | -97.72340 | United States | GBIF, 2025 |
| *D. maidis* | 41.75047 | -91.53995 | United States | GBIF, 2025 |
| *D. maidis* | 32.31177 | -97.01598 | United States | GBIF, 2025 |
| *D. maidis* | 33.03536 | -96.72633 | United States | GBIF, 2025 |
| *D. maidis* | 30.36642 | -97.67348 | United States | GBIF, 2025 |
| *D. maidis* | 31.63464 | -98.24814 | United States | GBIF, 2025 |
| *D. maidis* | 31.63464 | -98.24814 | United States | GBIF, 2025 |
| *D. maidis* | 18.60095 | -98.84515 | México | GBIF, 2025 |
| *D. maidis* | 18.60095 | -98.84515 | México | GBIF, 2025 |
| *D. maidis* | 40.00700 | -83.03000 | United States | GBIF, 2025 |
| *D. maidis* | 45.3999 | -72.8518 | Canada | Jacques et al. 2025 (in preparation) |
| *D. maidis* | 45.5868 | -73.0612 | Canada | Jacques et al. 2025 (in preparation) |
| *D. maidis* | 46.3976 | -72.7646 | Canada | Jacques et al. 2025 (in preparation) |
| *D. maidis* | 46.4552 | -71.7505 | Canada | Jacques et al. 2025 (in preparation) |
| *D. maidis* | 45.5117 | -72.9778 | Canada | Jacques et al. 2025 (in preparation) |
| *D. maidis* | 42.2092 | -93.4244 | United States | Current study |
| *D. maidis* | 36.13339 | -97.10624 | United States | Current study |
| *D. maidis* | 35.03601 | -97.90017 | United States | Current study |
| *D. maidis* | 37.01117 | -93.59683 | United States | Current study |
| *D. maidis* | 38.90684 | -92.28218 | United States | Current study |
| *D. elimatus* | 18.7149 | -97.3408 | México | GBIF, 2025 |
| *D. elimatus* | 19.8333 | -99.9833 | México | GBIF, 2025 |
| *D. elimatus* | 19.3000 | -98.7333 | México | GBIF, 2025 |
| *D. elimatus* | 19.8333 | -99.9833 | México | GBIF, 2025 |
| *D. elimatus* | 21.5333 | -101.7167 | México | GBIF, 2025 |
| *D. elimatus* | 20.0500 | -98.5333 | México | GBIF, 2025 |
| *D. elimatus* | 19.3000 | -98.7333 | México | GBIF, 2025 |
| *D. elimatus* | 19.8333 | -99.9833 | México | GBIF, 2025 |
| *D. elimatus* | 20.1667 | -98.3000 | México | GBIF, 2025 |
| *D. elimatus* | 19.8333 | -99.9833 | México | GBIF, 2025 |
| *D. elimatus* | 20.1667 | -98.3000 | México | GBIF, 2025 |
| *D. elimatus* | 19.8333 | -99.9833 | México | GBIF, 2025 |
| *D. elimatus* | 19.3000 | -98.7333 | México | GBIF, 2025 |
| *D. elimatus* | 19.8081 | -101.6965 | México | GBIF, 2025 |
| *D. elimatus* | 19.8079 | -101.6961 | México | GBIF, 2025 |
| *D. elimatus* | 18.6009 | -98.8451 | México | GBIF, 2025 |
| *D. elimatus* | 18.6009 | -98.8451 | México | GBIF, 2025 |
| *D. elimatus* | 19.7943 | -104.2225 | México | Moya-Raygoza et al 2014 |
| *D. elimatus* | 21.2336 | -103.5003 | México | Moya-Raygoza et al 2014 |
| *D. elimatus* | 19.6073 | -98.9254 | México | Nault and Madden, 1985 |
| *D. elimatus* | 21.4751 | -103.0929 | México | Moya-Raygoza, 2002 |
| *D. elimatus* | 16.3635 | -96.6379 | México | Moya-Raygoza, 2002 |
| *D. elimatus* | 17.8517 | -96.2151 | México | Moya-Raygoza, 2002 |
| *D. elimatus* | 18.4848 | -97.4594 | México | Moya-Raygoza, 2002 |
| *D. elimatus* | 17.2465 | -96.8607 | México | Moya-Raygoza, 2002 |
| *D. elimatus* | 17.4531 | -97.2368 | México | Moya-Raygoza, 2002 |
| *D. elimatus* | 19.6761 | -103.9343 | México | Moya-Raygoza, 2002 |
| *D. elimatus* | 20.3270 | -102.0474 | México | Moya-Raygoza, 2002 |
| *D. elimatus* | 20.6488 | -101.3146 | México | Moya-Raygoza, 2002 |
| *D. elimatus* | 20.5567 | -100.6898 | México | Moya-Raygoza, 2002 |
| *D. elimatus* | 20.2319 | -98.9158 | México | Moya-Raygoza, 2002 |
| *D. elimatus* | 19.8779 | -100.4407 | México | Moya-Raygoza, 2002 |
| *D. elimatus* | 20.2207 | -99.2084 | México | Moya-Raygoza, 2002 |
| *D. elimatus* | 19.8719 | -103.5955 | México | Moya-Raygoza, 2007 |
| *D. elimatus* | 31.3629 | -109.6696 | United States | Triplehorn and Nault, 1985 |
| *D. elimatus* | 31.4682 | -108.5139 | United States | Triplehorn and Nault, 1985 |
| *D. elimatus* | 19.4923 | -98.8775 | Mexico | Maramorosch, 1959 |
| *D. elimatus* | 19.6218 | -98.9146 | Mexico | Heady and Nault, 1985 |
| MBS | -23.1669 | -48.0503 | Brazil | Orlovskis et al. 2017 |
| MBS | -20.5336 | -48.51694 | Brazil | Orlovskis et al. 2017 |
| MBS | -19.7669 | -44.40028 | Brazil | Orlovskis et al. 2017 |
| MBS | 18.6797 | -99.1164 | Mexico | Moya-Raygoza and Nault, 1998 |
| MBS | 19.5287 | -98.8484 | Mexico | Moya-Raygoza and Nault, 1998 |
| MBS | 20.4901 | -97.5463 | Mexico | Moya-Raygoza and Nault, 1998 |
| MBS | 19.7329 | -97.5754 | Mexico | Perez-Lopez et al. 2016 |
| MBS | 19.7586 | -97.5509 | Mexico | Perez-Lopez et al. 2016 |
| MBS | 19.7323 | -97.5970 | Mexico | Perez-Lopez et al. 2016 |
| MBS | 19.8930 | -97.5874 | Mexico | Perez-Lopez et al. 2016 |
| MBS | 19.6455 | -97.1047 | Mexico | Perez-Lopez et al. 2016 |
| MBS | -12.0174 | -75.2889 | Peru | Gamarra et al. 2022 |
| MBS | -11.9937 | -75.2340 | Peru | Gamarra et al. 2022 |
| MBS | -11.9712 | -75.2614 | Peru | Gamarra et al. 2022 |
| MBS | -11.9562 | -75.2911 | Peru | Gamarra et al. 2022 |
| MBS | -12.1206 | -75.2537 | Peru | Gamarra et al. 2022 |
| MBS | -12.0982 | -75.2905 | Peru | Gamarra et al. 2022 |
| MBS | -12.079 | -75.3254 | Peru | Gamarra et al. 2022 |
| MBS | -12.0493 | -75.2902 | Peru | Gamarra et al. 2022 |
| MBS | -11.8344 | -75.4050 | Peru | Gamarra et al. 2022 |
| MBS | -10.4923 | -48.3144 | Brazil | Toloy et al. 2024 |
| MBS | -12.2160 | -46.0381 | Brazil | Galvao et al. 2020 |
| MBS | -17.8504 | -50.9295 | Brazil | Galvao et al. 2020 |
| MBS | -18.0182 | -49.3846 | Brazil | Galvao et al. 2020 |
| MBS | -17.6275 | -47.7885 | Brazil | Galvao et al. 2020 |
| MBS | -15.8446 | -43.3385 | Brazil | Galvao et al. 2020 |
| MBS | -18.9711 | -46.9784 | Brazil | Galvao et al. 2020 |
| MBS | -21.4844 | -47.0473 | Brazil | Galvao et al. 2020 |
| MBS | -21.7788 | -47.1220 | Brazil | Galvao et al. 2020 |
| MBS | -23.4427 | -48.9035 | Brazil | Galvao et al. 2020 |
| MBS | -28.6303 | -53.6330 | Brazil | Stürmer et al. 2024 |
| MBS | -20.0322 | -45.9868 | Brazil | Da Cunha et al. 2023 |
| MBS | -26.1874 | -50.4264 | Brazil | Canale et al. 2024 |
| MBS | -22.6906 | -50.4770 | Brazil | Gomes et al. 2004 |
| MBS | -24.0225 | -48.3687 | Brazil | Gomes et al. 2004 |
| MBS | -22.3101 | -54.8318 | Brazil | Gomes et al. 2004 |
| MBS | -21.0237 | -47.8078 | Brazil | Gomes et al. 2004 |
| MBS | 26.4341 | -98.2210 | United States | Harrison et al. 1996 |
| MBS | 25.4227 | -80.5491 | United States | Harrison et al. 1996 |
| MBS | 10.6634 | -85.5146 | Costa Rica | Harrison et al. 1996 |
| MBS | 20.5805 | -97.4575 | Mexico | Harrison et al. 1996 |
| MBS | 18.6831 | -99.1284 | Mexico | Harrison et al. 1996 |
| MBS | 4.692313 | -75.963432 | Colombia | Dr. Javier Orozco, Alphanalitica S.A.S, 2022 |
| MBS | 3.855608 | -76.308398 | Colombia | Dr. Javier Orozco, Alphanalitica S.A.S, 2022 |
| MBS | 3.348848 | -76.228093 | Colombia | Dr. Javier Orozco, Alphanalitica S.A.S, 2022 |
| MBS | 2.668382 | -75.333302 | Colombia | Reinoso, 2021 |
| MBS | 2.869621 | -75.281099 | Colombia | Reinoso, 2021 |
| MBS | 2.225862 | -75.611987 | Colombia | Reinoso, 2021 |
| MBS | 2.108519 | -75.831831 | Colombia | Reinoso, 2021 |
| MBS | 42.2092 | -93.4244 | United States | Current study |
| MBS | 36.13339 | -97.10624 | United States | Current study |
| MBS | 35.03601 | -97.90017 | United States | Current study |
